# CD33-CD45 Interaction Reveals a Mechanistic Link to Alzheimer’s Disease Susceptibility

**DOI:** 10.1101/2025.07.28.667311

**Authors:** Nicole Vo, Cory D. Rillahan, Zena K. Chatila, Daniel M. Virga, Jennifer L. Hall, Kirstin A. Tamucci, Mamunur Rashid, Sarah M. Connor, Sana Chintamen, Gaelen Guzman, Mariko Taga, Minghua Liu, Philip L. De Jager, Peter St. George-Hyslop, Annie J. Lee, Badri N. Vardarajan, David A. Bennett, Jennifer J. Kohler, Monica Schenone, Wassim Elyaman, Steven A. Carr, Elizabeth M. Bradshaw

**Author notes:** Correspondence should be addressed to E.M.B. Equal contribution (Nicole Vo, Cory D. Rillahan, Zena K. Chatila, Daniel M. Virga).

## Abstract

The innate immune gene *CD*33, encoding a myeloid inhibitory sialic acid-binding receptor, is associated with Alzheimer’s disease (AD) susceptibility. The AD-associated *rs3865444^CC^* risk variant reduces splicing of the sialic acid-binding domain and increases expression of the full-length (sialic acid-binding) CD33 isoform seven-fold compared to the *rs3865444^AA^* protective genotype. Here we identify CD45 as an immune cell-specific sialic acid-dependent *cis* CD33 binding partner, whose phosphatase activity is inhibited by CD33. Overexpression of CD33 or loss of CD45 contributes to impaired microglial clearance of amyloid beta and amyloid beta-induced loss of dendritic spines in microglial-neuronal co-cultures, aligning with a detrimental effect of CD33-mediated inhibition of CD45. CD33-CD45 interaction frequency was increased in monocytes from individuals with the *rs3865444^CC^* risk variant compared to *rs3865444^AA^*, as well as in AD compared to controls, independent of genotype. Furthermore, an interaction between *CD33* and *PTPRC* (encoding CD45) gene expression in human brain tissue was associated with a pathological diagnosis of AD and global burden of AD pathology. Our findings thus establish a functional interaction between CD33 and CD45 relevant to AD susceptibility and systemic myeloid dysfunction in this disease.

## Introduction

CD33 is a sialic acid binding inhibitory transmembrane receptor predominantly expressed on cells of the myeloid lineage^1^, although it can be upregulated on a subset of adaptive immune cells^2^. In the central nervous system (CNS), it is highly expressed in microglia, the resident innate-immune myeloid cells^3^. Microglia not only protect the CNS from foreign infections but also help maintain the health of neurons using processes that require a fine balance of inhibitory and activating signaling to carry out their requisite functions^4, 5^. To this end, microglia express many cell surface recognition molecules that tie extracellular events to intracellular signaling. Among these are a variety of lectins including Siglecs (sialic acid-binding immunoglobulin-like lectins), which are thought to regulate the innate immune inflammatory response^6^. CD33, also known as Siglec-3, is thought to be a negative regulator of microglial function as it contains an immunoreceptor tyrosine-based inhibitory motif (ITIM) which interacts with the phosphatase SHP-1 and has been shown to constitutively repress pro-inflammatory cytokines in monocytes^7, 8^.

Genome-wide association studies (GWAS) have identified the CD33 locus as a risk factor for AD^9, 10^. We previously demonstrated that individuals with the AD-associated *rs3865444^CC^* risk allele have increased expression of the full-length isoform of CD33 (CD33^M^) on the surface of their innate immune cells compared to those with the *rs3865444^AA^* protective genotype^11–13^. The risk allele is also associated with 1) diminished internalization of amyloid-β1-42 peptide (Aβ_1-42_), 2) accumulation of neuritic amyloid pathology in the brain, 3) higher fibrillar amyloid levels on *in vivo* positron emission tomography imaging, and 4) increased numbers of human microglia with small, thick processes and a rounded morphology^11^. The causative single nucleotide polymorphism (SNP) underlying the association between *CD33* and AD is most likely *rs12459419*, which is in high linkage disequilibrium with the SNP *rs3865444* that was identified through GWAS^12, 14^. The SNP *rs12459419* is thought to modulate splicing of exon 2 of CD33, which encodes the sialic acid-binding domain, resulting in increased expression of the exon 2 containing isoform of CD33 (CD33^M^) in individuals with the risk variant. These findings suggest that the sialic acid-binding domain is critical for the genetic association of the *CD33* locus with susceptibility to AD. CD33 is known to bind α2,6-linked and α2,3-linked sialic acid bearing glycans^15, 16^. However, the protein binding partners carrying these molecules, and thereby regulated by CD33 in myeloid cells, are currently unknown. Here, we identify myeloid sialic acid-specific binding partners of CD33, including CD45. We demonstrate that CD33 suppresses CD45 phosphatase activity, and that the interaction between CD33 and CD45 is modified by *CD33* genotype and AD disease status. We further investigate the functional implications of this interaction in microglia in the context of amyloid pathology.

## Results

### Identifying novel CD33 binding partners specific to the AD-associated sialic acid binding domain

Since the Alzheimer’s disease risk allele at the CD33 locus leads to increased expression of the CD33 isoform containing the sialic acid binding domain, we hypothesized that to understand the role of CD33 in AD, we must determine the CD33 sialic acid-specific binding partners in human myeloid cells. To identify these binding partners, we incubated the human myeloid cell line THP-1 with a mannosamine analog, Ac_4_ManNDAz^17, 18^. This molecule is taken up by cells through passive diffusion, deacetylated, and converted into the corresponding sialic acid containing a photocrosslinker at a position that does not interfere with CD33 binding. When treated with UV light, a covalent connection is made between the modified sialic acid and neighboring molecules, allowing for the *in situ* identification of the glycoproteins interacting with CD33 by subsequent immunoprecipitation (IP) and nanoscale liquid chromatography coupled to mass spectrometry (nanoLC-MS). This crosslinking strategy allowed us to investigate protein-glycan interactions, which (i) may not survive the stringent washing conditions of the immunoprecipitation process and (ii) can be dictated by membrane co-localization in micro domains (i.e. lipid rafts) which are destroyed upon cell lysis^19^.

In comparison to control THP-1 cells either not given the photocrosslinking sugar or not treated with UV light, we identified CD43, CD45, GLUT1, and CLEC11a as myeloid expressed sialic acid-specific binding partners for CD33 with adjusted p-values of less than 0.05 (Figure 1a, 1b, Supplementary Data Tables 1 and 2, and Extended Data Figure 1). CLEC11a was also increased without crosslinking compared to IgG, suggesting that it is not dependent on the sialic acid-specific interaction, but rather enhanced.

**Figure 1:**
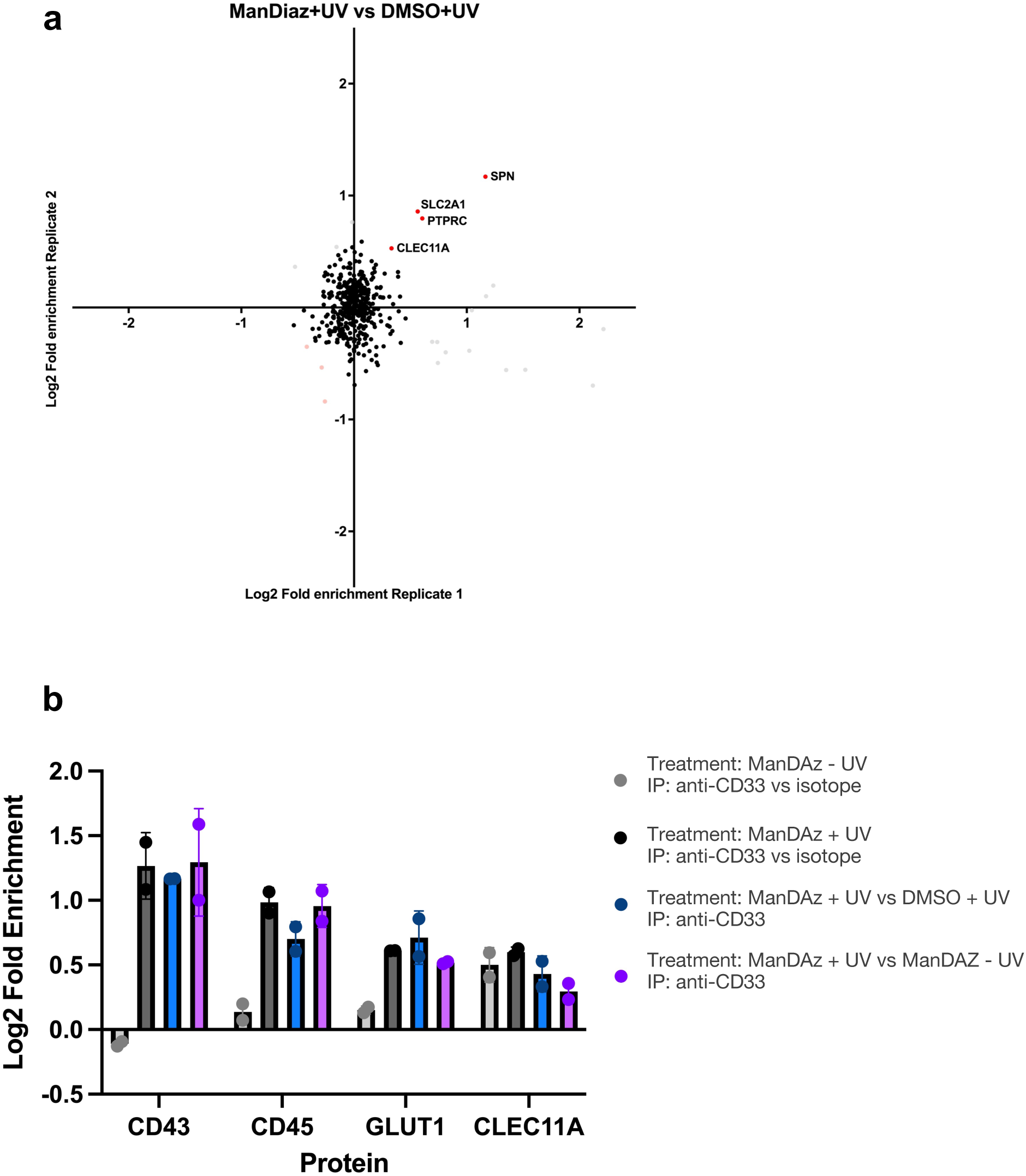
NanoLC-MS identifies CD43, CD45, GLUT1, and CLEC11A as sialic acid-specific binding partners of CD33. **a.** Proteins that were significantly enriched as CD33 binding proteins, defined as associated with CD33 after THP-1 cells were treated with Ac_4_ManNDAz and treated with UV light compared to THP-1 cells treated with DMSO alone and UV light. **b.** Proteins that were not enriched in the CD33 immunoprecipitation without Ac_4_ManNDAz compared to IgG control immunoprecipitation, demonstrating their specificity for the sialic acid binding domain. These proteins were enriched after the Ac_4_ManNDAz plus UV light treatment compared to the Ac_4_ManNDAz treatment without UV light crosslinking. Error bars denote the standard deviation between technical replicates.

### CD33 and CD45 binding occurs in *cis* in a sialic acid-dependent manner and is regulated by the *CD33 rs3865444* genotype

As loss of CD45, an immune cell specific phosphatase, has been implicated in AD, we further investigated whether its interaction with CD33 has functional consequences that may contribute to AD risk^20–23^. We first validated CD45-CD33 binding in THP-1 cells utilizing an orthogonal approach, the proximity ligation assay (PLA), which sensitively identifies proteins in close proximity (40 nm or less) without a need for a crosslinking strategy (Figure 2a). When CD33 protein expression was eliminated with CRISPR/Cas9, the PLA signal between CD33 and CD45 was no longer present, confirming PLA signal was due to CD33-CD45 binding (Figure 2a and Extended Data Figure 2). We also confirmed that this interaction between CD33 and CD45 occurs in *cis*. We combined THP-1 cells with either CD33 or CD45 knocked out (KO) by CRISPR/Cas9, so that no cell expressed both molecules. We found that the interaction between the two molecules was disrupted when the two molecules were not expressed on the same cell, demonstrating that this interaction occurs in *cis* and is not observed in *trans* (Figure 2b, Extended Data Figure 2, Extended Data Figure 3)^24^. We further validated this interaction using a solid phase binding assay. Using two concentrations of human CD45 produced in a mouse myeloma cell line to maintain glycosylation, we confirmed that the extracellular domains of CD33 and CD45 directly bind (Figure 2c). This CD33-CD45 binding was disrupted upon the addition of sialidase (Extended Data Figure 4). These findings validate that the extracellular domains of CD33 and glycosylated CD45 interact in a sialic acid-dependent manner.

**Figure 2:**
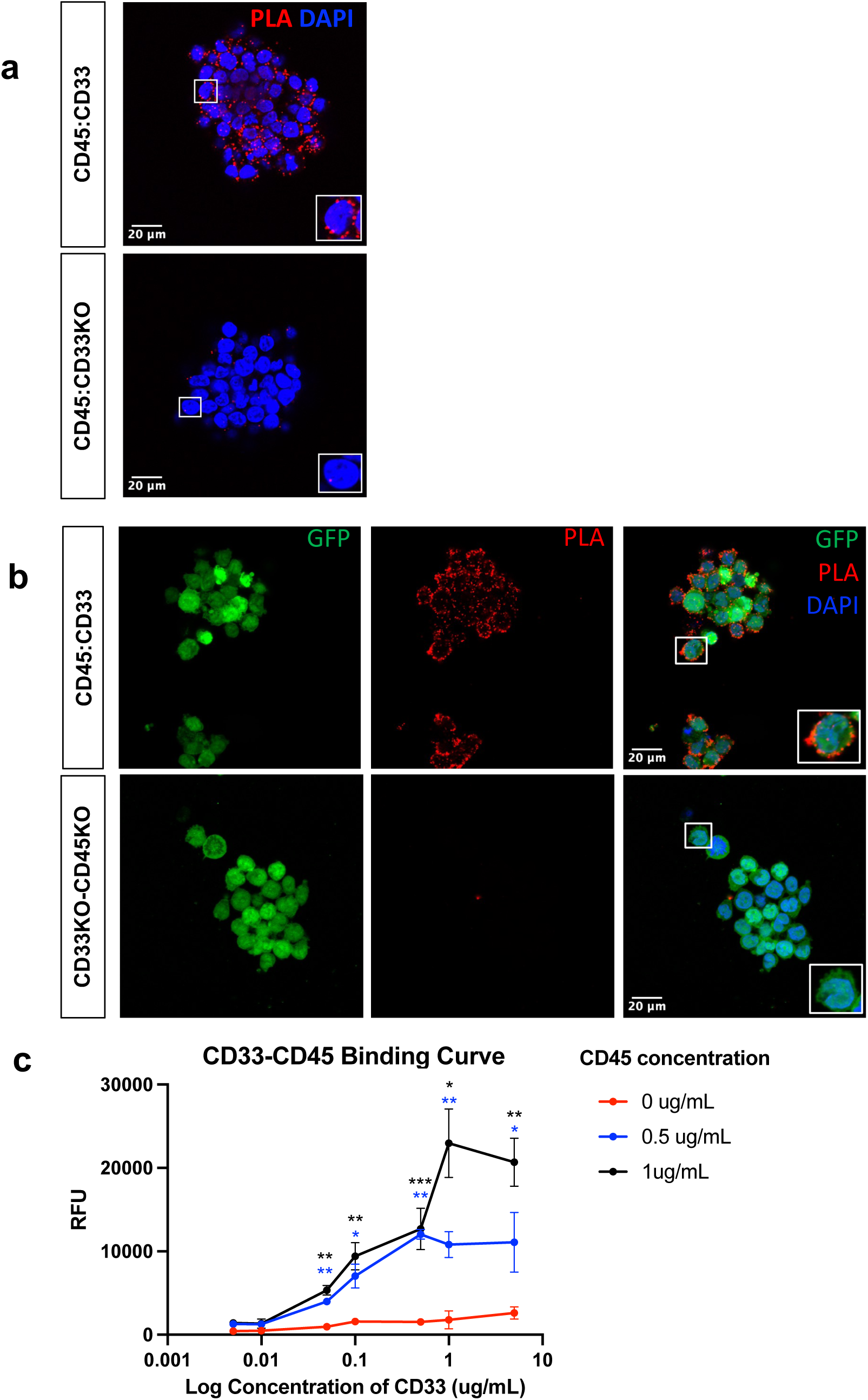
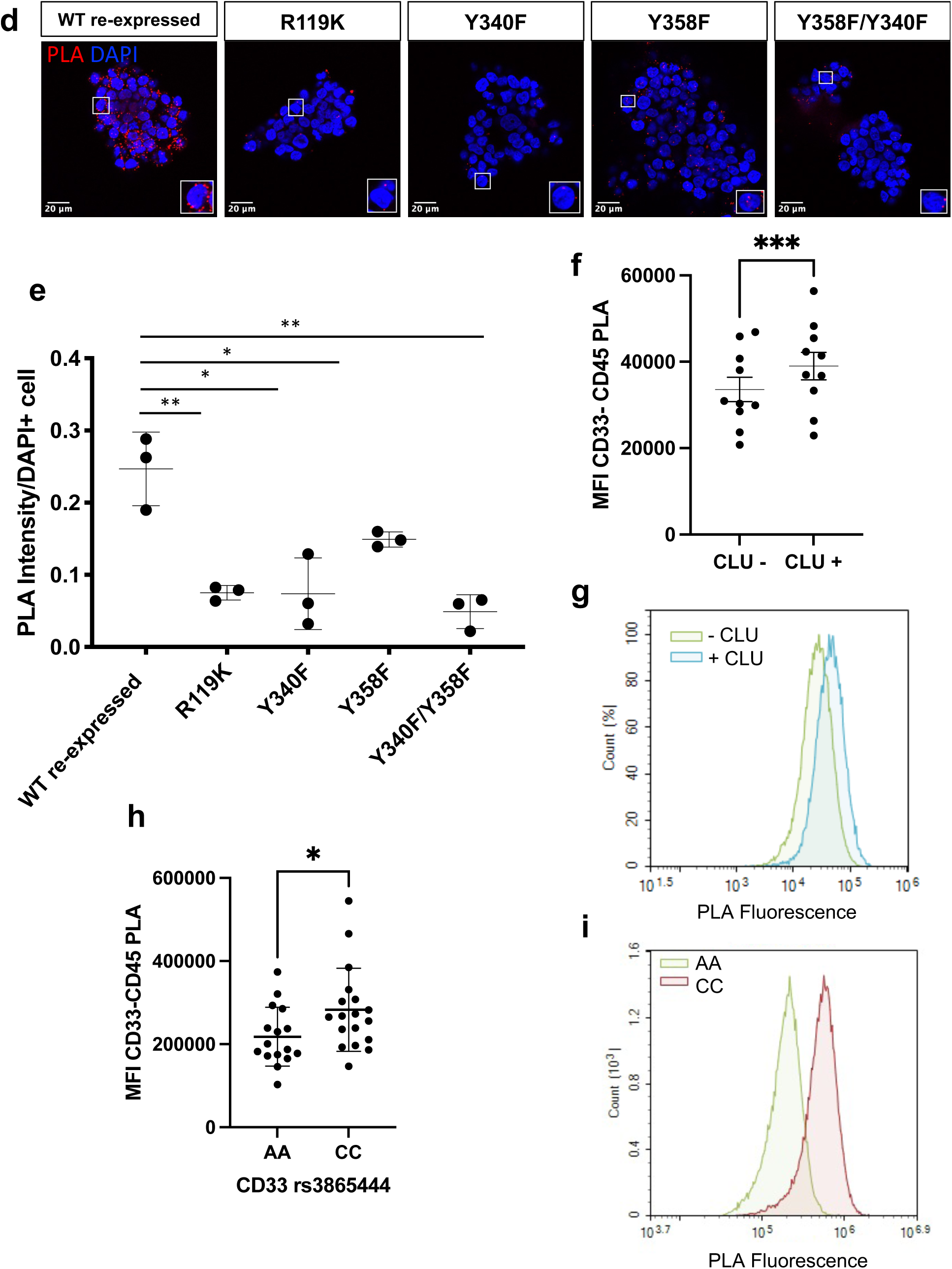
CD33 binds to CD45 in *cis* in a *CD33 rs3865444* genotype-dependent manner. **a.** The mass spectrometry findings of the CD33 and CD45 interaction were validated in THP-1 cells using the proximity ligation assay (PLA). Knock-out (KO) of CD33 from the THP-1 cell line led to the elimination of the interaction between CD33 and CD45, as detected by PLA. **b.** Combining THP-1 cells with either CD33 KO or CD45 KO via CRISPR/Cas9 leads to a reduction in the interaction between CD33 and CD45, suggesting that the interaction is occurring in *cis.* **c.** In a solid phase binding assay, recombinant human CD33 bound to immobilized recombinant human CD45 in a dose dependent manner. Increasing concentrations of recombinant hCD33-Fc (0.005µg/mL-5µg/mL) were incubated overnight at 4°C to recombinant hCD45 coated plates and an anti-human Fc antibody was used to detect recombinant bound hCD33. Each dot represents the average of three independent experiments. Two-way ANOVA with Tukey’s multiple comparison test, p*<0.05, p**<0.01, p***<0.001. Asterisks denote comparison of 0.5 μg/mL (blue asterisks) or 1 μg/mL (black asterisk) to 0 μg/mL CD45 concentration treatment. **d.** Representative images of CD33-CD45 PLA signal in CD33 mutant THP-1 cell lines. Mutations in the sialic acid binding domain (R119K) and the ITIM (Y340F) disrupt the interaction between CD33 and CD45 in THP-1 monocytes, as demonstrated by reduced PLA signal. Mutation in the ITIM-like domain (Y358F) leads to a reduction in CD33-CD45 interaction, as does the double mutant (Y358F/Y340F). **e.** Quantification of the CD33-CD45 PLA signal in each mutant THP-1 line, as PLA intensity per DAPI positive cell. Each dot represents a technical replicate. Unpaired t-test. *p<0.05, **p<0.01. **f.** Median fluorescence intensity of CD33-CD45 Flow Cytometry PLA in primary monocytes from 10 individuals, with and without pretreatment with 60nM clusterin (CLU) for 15 minutes. Error bars = mean ± SEM. ***p<0.001, paired t-test. **g.** Representative overlay flow cytometry histogram of CD33-CD45 PLA fluorescent signal from monocytes with and without pretreatment with 60nM clusterin for 15 minutes. **h.** Median fluorescence intensity of CD33-CD45 PLA in primary monocytes from individuals genotyped for *CD33 rs3865444*. N = 16 *rs3865444^AA^* and 18 *rs3865444^CC^.* Error bars = mean ± SEM. *p<0.05, unpaired t-test. **i.** Representative overlayed flow cytometry histogram of PLA fluorescent signal from one *CD33 rs3865444^AA^* and one *CD33 rs3865444^CC^* individual.

In order to further validate that the CD33 and CD45 interaction is dependent on the sialic acid-binding domain of CD33, we expressed CD33 containing a mutation at the critical sialic acid binding amino acid arginine 119 (R119)^25^ in CD33-deleted THP-1 cells and performed PLA. No signal was detected, indicating that mutating CD33 at R119 eliminated the interaction between CD33 and CD45 when compared to the re-expression of wild-type CD33^M^ (Figure 2d, 2e, Extended Data Figure 5). This finding further confirmed not only that CD33 and CD45 interact, but that the interaction is sialic acid-dependent.

To examine whether, in addition to the sialic acid-dependent extracellular interaction, there is an intracellular interaction between CD33^M^ and CD45, we mutated the key tyrosine residues in the ITIM and ITIM-like motifs of CD33^M^ and again measured CD33 and CD45 interaction using PLA. Tyrosine 340 (Y340) in the ITIM motif is known to be required for the binding of SHP-1, a phosphatase often utilized by CD33 for intracellular signaling that we demonstrated interacts with CD33 in a *rs3865444* genotype-dependent manner^8, 25^. Interestingly, mutating Y340 led to a complete disruption of the interaction between CD33^M^ and CD45, similar to our findings with the R119K sialic acid binding domain mutation (Figure 2d, 2e). Mutating tyrosine 358 (Y358) in the ITIM-like motif reduced the interaction, but not to the same extent as R119K. As expected, the double mutant Y340/Y358 also significantly reduced the interaction. These findings indicate that the interaction between CD33^M^ and CD45 requires both extracellular binding at the sialic acid binding domain and CD33 signaling through its ITIM.

While the accompanying manuscript (*Dodd, et al.*) demonstrates that CD33 is a dimer, little is known about how the binding of one sialylated partner may affect binding of other partners to CD33. To investigate how *trans* binding partners, such as soluble clusterin (identified in the accompanying manuscript), impact CD45 binding to CD33, we performed PLA investigating the CD33-CD45 interaction on monocytes with and without the addition of clusterin. Interestingly, we found that the addition of clusterin increased the interaction between CD33 and CD45 (Figure 2f, 2g). This finding may be due to complex binding dynamics between ligands and the CD33 dimer, such as extracellular CD33 ligand binding enhancing the binding ability of the dimerized partner. Alternatively, extracellular engagement of the CD33 sialic acid binding domain may enhance intracellular interactions with other molecules via the ITIM motif.

We have previously shown that the AD-associated SNP *rs3865444* leads to increased expression of the sialic acid-binding domain containing isoform of CD33 (CD33^M^) on the surface of monocytes and monocyte-derived microglia-like cells (MDMi), an *in vitro* primary human microglial model^11, 13, 26^. To investigate whether the *CD33* genotype impacts the CD33-CD45 interaction, we examined the interaction of these two molecules by PLA in *ex vivo* monocytes from genotyped individuals. We found that in the primary monocytes, as in the THP-1 cells, CD33 and CD45 are in close enough proximity to be interacting by PLA, validating the interaction we observed in THP-1 cells. Monocytes from individuals with the AD-associated risk *rs3865444^CC^* genotype had significantly more interactions between CD33 and CD45 than monocytes from individuals with the protective SNP, *rs3865444^AA^* (Figure 2h, 2i, Extended Data Figure 6). This finding is consistent with reduced CD33^M^ expression on the surface of monocytes from individuals with the protective genotype^11^. To confirm this interaction on primary human microglia, we next examined the interaction *in situ* in post-mortem human brain tissue (Supplementary Data Table 3). We found that CD45-positive cells within the cortex of the CNS, predominantly microglia, were positive by PLA for interactions between CD33 and CD45 (Extended Data Figure 7). Upon quantifying the interaction, we found that CD45-positive cells from individuals with the AD-associated risk *rs3865444^CC^* genotype had more interactions between CD33 and CD45 than microglia from individuals with the protective SNP, *rs3865444^AA^* (Extended Data Figure 7). These findings suggest that there is increased interaction between CD33 and CD45 in both the peripheral immune system and CNS of individuals with the *CD33* risk variant.

### CD33 inhibits CD45 phosphatase activity

Siglecs are known to act as negative regulators of their binding partners and thereby fine-tune cell signaling events^7^. This led us to investigate whether CD33 similarly negatively regulates CD45 function, specifically its phosphatase activity. Using THP-1 WT and THP-1 CD33KO cells, we first examined the effect of CD33 on global phosphatase activity. We found that CD33 KO led to an increase in phosphatase activity, both with and without LPS activation, to provide an inflammatory stimulus (Figure 3a). To determine if this increase in phosphatase activity was due to the ability of CD33 to bind to sialic acid, we compared THP-1 CD33KO cells with re-expressed wild-type CD33^M^ or the R119K mutant. We found that both at baseline and in the presence of LPS stimulation, cells expressing the R119K mutant version of CD33, which cannot bind to sialic acid, have greater phosphatase activity (Figure 3b), supporting that CD33 binding via its sialic acid binding domain reduces phosphatase activity.

**Figure 3:**
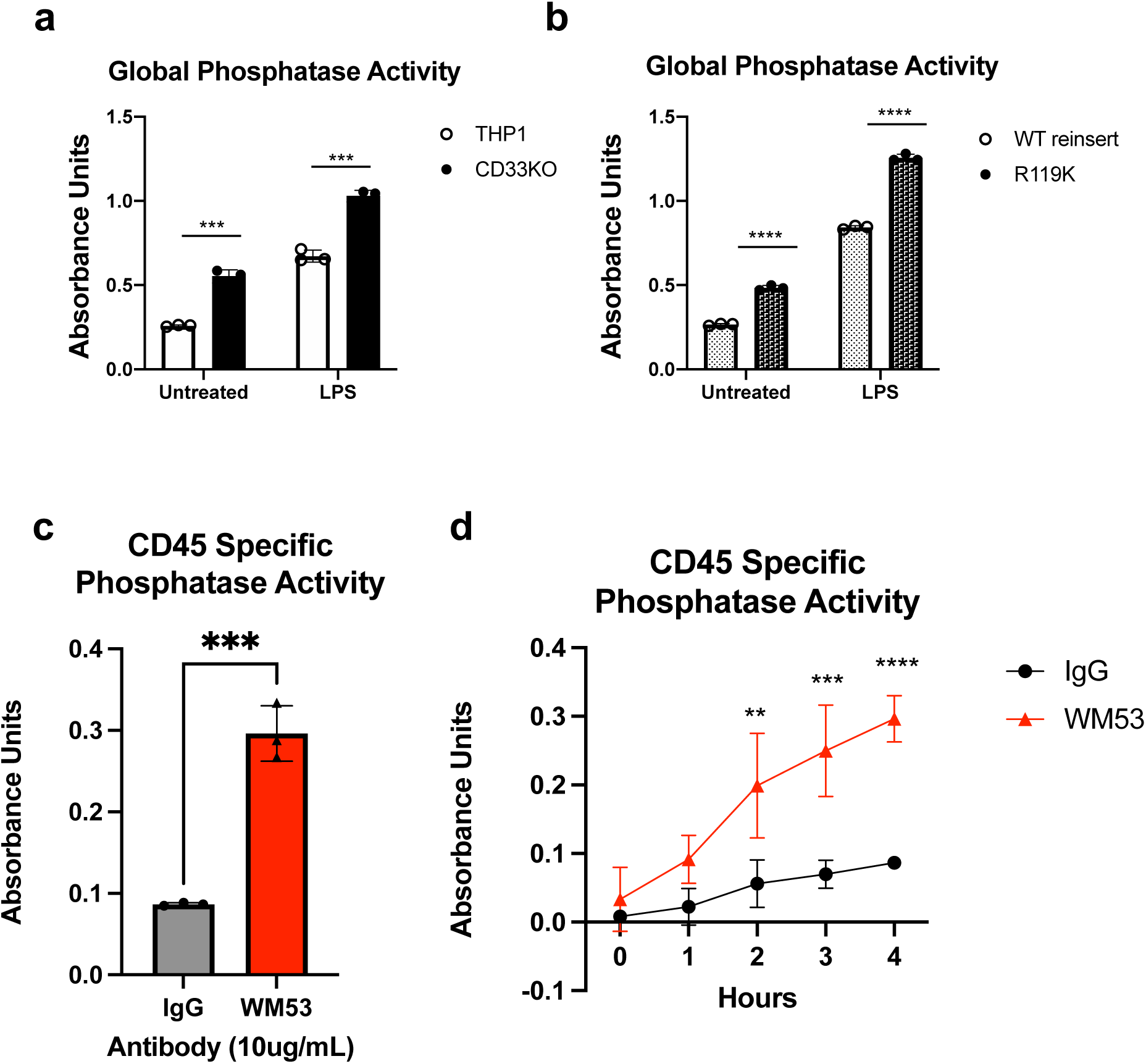
CD33 Inhibits CD45 Phosphatase Activity. **a.** CD33KO THP-1 cells have higher phosphatase activity compared to WT THP-1 cells, both at baseline and after stimulation with LPS. Phosphatase activity was measured with a colorimetric assay, reported with absorbance units. Unpaired t-test. ***p<0.001. **b.** Mutant CD33KO THP-1 monocytes with CD33 R119K re-expressed have higher phosphatase activity compared to CD33KO THP-1 cells with WT CD33 re-expressed, both at baseline and after stimulation with LPS. Unpaired t-test. ***p<0.001; ****p<0.0001. **c.** Treating THP-1 cells with the anti-CD33 antibody WM53, which profoundly reduces cell surface expression of CD33, increases CD45-specific phosphatase activity. CD45-specific phosphatase activity was determined by subtracting phosphatase activity measured with the addition of a CD45 specific inhibitor (CD45 Inhibitor VI), capturing all non-CD45 phosphatase activity, from global phosphatase activity. WM53 treatment was compared to IgG control. Each dot represents the average of two technical replicates. Unpaired t-test. ***p<0.001. **d.** CD45-specific phosphatase activity was measured over a period of 4 hours as described above. WM53 treatment of THP-1 monocytes, which profoundly reduces CD33 surface expression, maintains a consistent increase in CD45-specific phosphatase activity compared to IgG control over the course of the reaction. Each dot represents the mean ± SEM of three experiments for WM53 and IgG, and each experiment was the average of two technical replicates. Two-way ANOVA with Sidak’s multiple comparisons test, **p<0.01, ***p<0.001, ****p<0.0001.

We next investigated the impact of CD33 binding on CD45-specific phosphatase activity in THP-1s using a CD45 inhibitor. We downregulated CD33^M^ expression in THP-1s using antibody clone WM53, an antibody that dramatically reduces CD33 expression at the cell surface without impacting CD45 expression (Extended Data Figure 8)^13^. We then measured CD45-specific phosphatase activity by subtracting phosphatase activity measured with a CD45 inhibitor (CD45 Inhibitor VI), capturing all non-CD45-mediated phosphatase activity, from global phosphatase activity. Compared to an IgG control, CD33 downregulation with WM53 treatment increased CD45-specific phosphatase activity (Figure 3c, 3d). Together, these findings demonstrate that CD33 binding at the sialic acid binding domain specifically inhibits CD45 phosphatase activity.

### CD33 overexpression and CD45 inhibition reduce amyloid beta clearance and contribute to amyloid-mediated synapse loss

As CD33 is genetically associated with AD, and its interaction with CD45 is specific to the isoform increased with the AD-risk variant, we next explored this interaction in relation to Alzheimer’s disease pathology. Having established that CD33^M^ inhibits CD45 phosphatase activity, we next examined whether this interaction between CD33^M^ and CD45 changes in the context of amyloid pathology. First, we stressed MDMi, a primary monocyte-derived model of human microglia, with aggregated Aβ_1-42_ and found that exposure to Aβ_1-42_ increased the number of interactions between CD33^M^ and CD45 per cell (Figure 4a, 4b)^26^. We measured CD33 and CD45 protein expression and found that Aβ_1-42_ exposure increased expression of both molecules (Figure 4c, 4d), aligning with a previous report demonstrating that CD33 protein expression is increased on microglia in the AD brain^27^. Our findings suggest that increased CD33-CD45 interaction in the context of Aβ_1-42_ is due to the Aβ_1–42_–induced upregulation of CD33 and CD45 protein expression on the surface of the cells.

**Figure 4:**
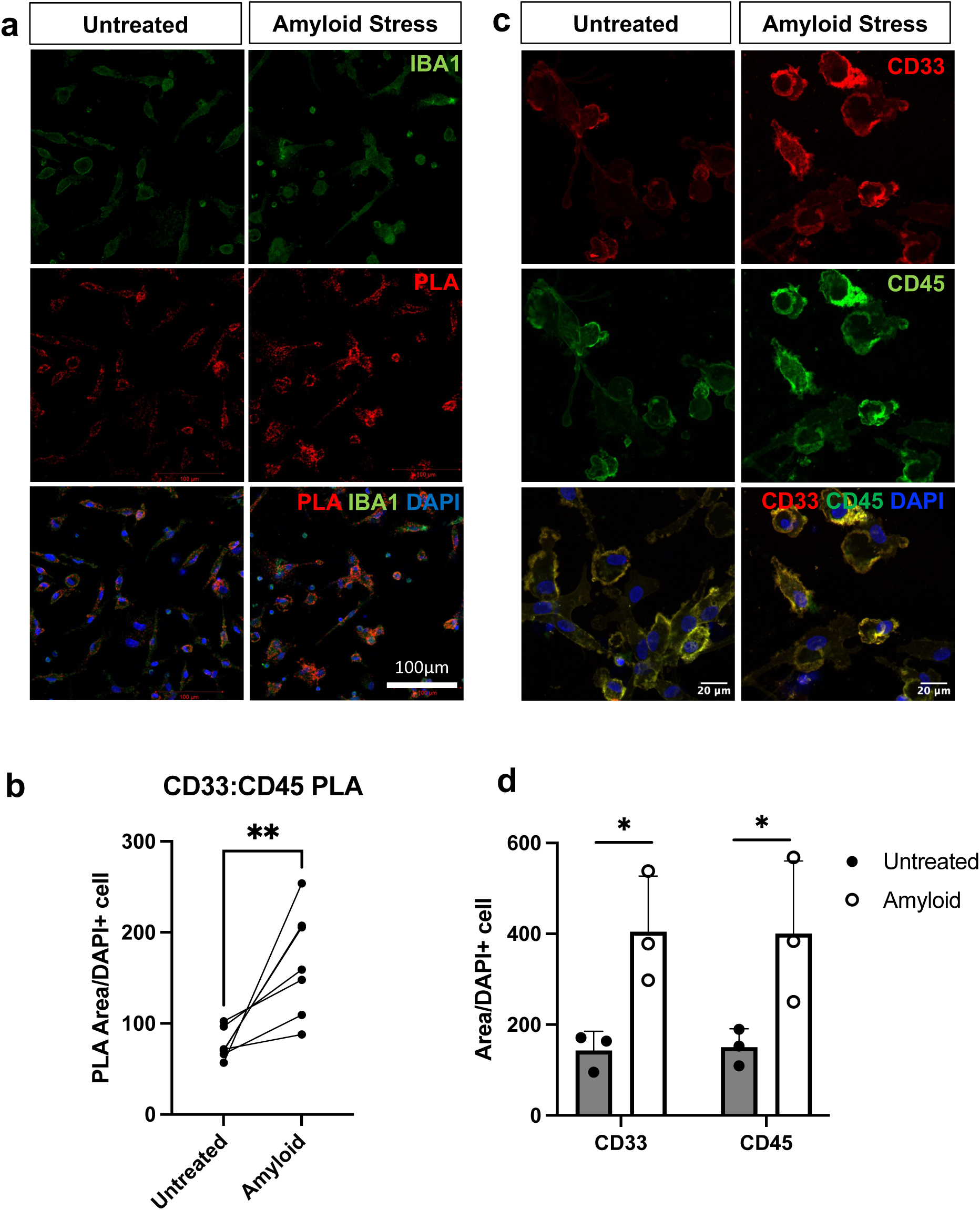
Amyloid stress increases the frequency of CD33-CD45 interaction in monocyte-derived microglia. **a.** Primary human monocyte-derived microglia (MDMi) were treated with aggregated Aβ_1-42_ for 24 hours, after which proximity ligation assay (PLA) was performed to examine CD33-CD45 interaction. Representative images of PLA signal for untreated and Aβ_1-42_ treated cells are shown. **b.** PLA signal was quantified as the PLA signal area per DAPI+ cell. Each dot represents an individual with paired untreated and Aβ_1-42_ treated cells. There was an increase in the number of interactions between CD33 and CD45 after aggregated Aβ_1-42_ stress compared to untreated MDMi from the same individuals. N=6 biological replicates. Ratio paired t-test. **p<0.01. **c.** Representative images of CD33 and CD45 MDMi protein staining following treatment with Aβ_1-42_ for 24 hours. **d.** CD33 and CD45 protein expression was quantified as area of fluorescent signal for each DAPI+ cell. N=3 biological replicates. Ratio paired t-test Aβ_1-42_ stress relative to untreated control. *p<0.05.

We previously linked the CD33 genotype to modulation of amyloid-beta uptake^11^ such that the risk variant leads to a reduction of uptake driven by the expression of CD33^M27^. However, it is unclear how this impaired phenotype impacts neuronal health and synaptic density, which has been strongly correlated with cognitive decline in AD^28–30^. Loss of synaptic density has been found to be an early event in AD^31, 32^, and microglia have been both positively and negatively implicated in this process^33–35^. Therefore, using the human microglial clone 3 (HMC3) cell line co-cultured with long-term murine cortical pyramidal neurons *ex utero* electroporated with tdTomato^36, 37^ (Figure 5a), we evaluated the relationship between microglial CD33^M^ and CD45 expression and Aβ_1-42_-mediated spine loss in neurons. Co-culturing HMC3 cells expressing GFP alone (plasmid control) in the presence of synaptically connected, fully developed pyramidal neurons was not sufficient to induce significant changes in spine density or neuronal morphology (Figure 5b, 5e). Upon the addition of oligomerized Aβ_1-42_, the neurons exhibited a significant reduction in dendritic spine density^38^. In addition to spine density, we measured Aβ_1-42_ uptake by the HMC3 cells in the co-culture. By imaging at high resolution, we volumetrically captured and quantified individual microglia as well as the intracellular fluorescent Aβ_1-42_ consumed by each microglial cell (Figure 5c, 5d). Not only did the loss of CD33 expression in HMC3 cells lead to an increase of Aβ_1-42_ uptake, but it also rescued amyloid-induced loss of dendritic spines (Figure 5b, 5d, 5e, and Extended Data Figures 9, 10, 11, 12). Conversely, overexpression of CD33^M^ led to a reduction in amyloid internalization but had no effect on amyloid-induced spine density reduction in primary neurons compared to the plasmid control. Knocking out CD45 expression resulted in the same phenotype as CD33^M^ overexpression (Figure 5b, 5d, 5e, and Extended Data Figures 9, 10, 11, 12), aligning with our findings that CD33^M^ inhibits CD45 phosphatase activity. Altogether, these results suggest that CD33^M^ activity and CD45 inhibition impair Aβ_1-42_ clearance and microglial protective functions, contributing to microglial amyloid-induced reduction of dendritic spines, aligning with our findings of CD33^M^ mediated inhibition of CD45.

**Figure 5.**
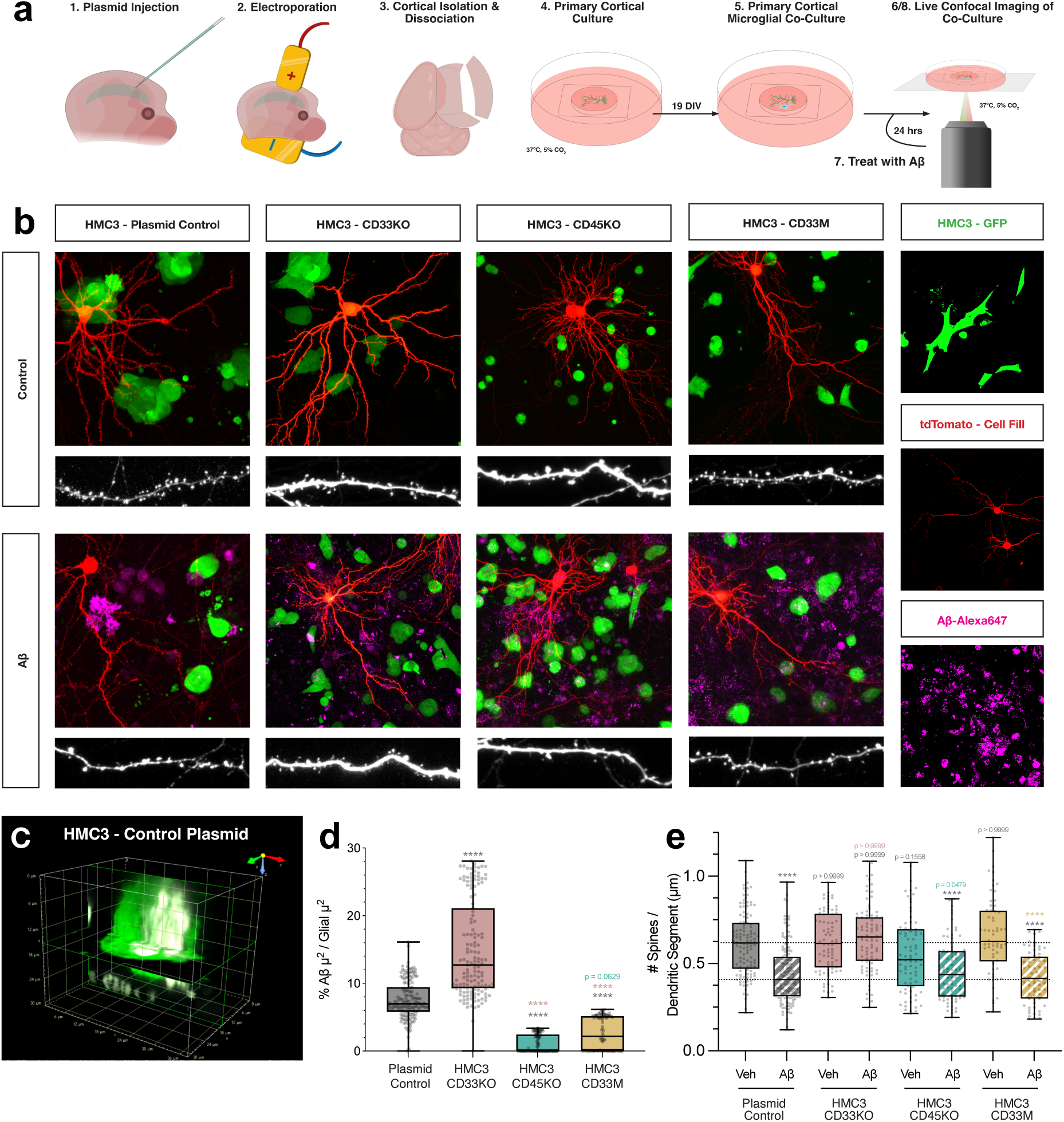
CD33KO prevents Aβ1-42-mediated spine loss and increases microglial Aβ1-42 consumption in primary neuronal and human microglial co-cultures. **a.** Schematic representation of the experimental paradigm. Briefly, E15.5 mouse embryos were electroporated with pCAG-tdTomato to target L2/3 cortical progenitors, The cortex is dissected, dissociated, and cultured for 18 days before human microglia carrying various alleles are added to the cultures. The cultures are then imaged live via confocal microscopy, treated with Aβ_1-42_ and imaged again. **b.** Representative images of sparsely electroporated primary cortical neurons (red, tdTomato) and human microglia (green, GFP) co-cultures before and after treatment with _Aβ1-42_ (magenta, Alexa647; 0.5 μg/ml) at 20 and 21DIV respectively. Additional insets include secondary dendritic segments and isolated channels. **c.** An x-y sliced, representative three-dimensional volumetric digital reconstruction of an individual HMC3-Control Plasmid microglia from neuronal-glial co-culture demonstrating Aβ_1-42_ consumption and co-localization. **d.** Quantification of Aβ_1-42_ consumption. All consumption analyses were done blind to experimental conditions as described in the materials and methods using NIS-Elements. Briefly, the microglial population’s total volume was quantified. The 3D area occupied by Aβ_1-42_ was then quantified and instances where the two signals co-localized, which is designated consumption, was divided by the total volume of the microglia as a function of percentage of microglia occupied by Aβ_1-42_. **e.** Quantification of spine density. All the spine density analyses were done blind to the experimental conditions and were done by manual counting using FIJI. All the statistical analyses were performed with Prism9 (GraphPad Software). Data is represented by box plots displaying Tukey error bars, with the box denoting 25th, 50th (median) and 75th percentile from 3-6 independent experiments. nControl Plasmid = 39 neurons, 117 dendritic segments; nControl Plasmid + Aβ = 49 neurons, 123 dendritic segments—6 biological replicates; nCD33KO = 27 neurons, 78 dendritic segments; nCD33KO + Aβ = 28 neurons, 83 dendritic segments— 3 biological replicates; nCD45KO = 27 neurons, 79 dendritic segments; nCD45KO + Aβ = 26 neurons, 78 dendritic segments—3 biological replicates; nCD33M = 19 neurons, 71 dendritic segments; nCD33M + Aβ = 22 neurons, 81 dendritic segments—3 biological replicates; Statistical analyses were performed using Kruskal-Wallis test followed by Dunn’s multiple comparisons. The tests were considered significant when p < 0.05. * p < 0.05; ** p < 0.01; *** p < 0.001; **** p < 0.0001; ns, not significant. Color of the p value indicates to what the condition is compared.

### CD33 and CD45 interaction increases in AD

To examine the implications of the interaction between CD33 and CD45 on AD, we examined the gene–gene interaction in a transcriptomics dataset of dorsolateral prefrontal cortex (DLPFC) from the ROSMAP cohort, a longitudinal cohort of aging^39^. To understand the biological significance of this interaction in the context of Alzheimer’s disease, we investigated whether the interaction between *CD33* and *PTPRC* (the gene which encodes CD45) gene expression influences AD clinical and neuropathological traits. After correction for multiple hypothesis testing, the *CD33-PTPRC* interaction was significantly associated with pathological diagnosis of AD and global AD pathology burden, both with negative beta values (Figure 6a). Their interaction was also nominally significant for tangles (Supplementary Data Table 4). The negative beta values of these associations indicate an inverse relationship between *CD33* and *PTPRC* gene expression in relation to pathology, aligning with our protein findings demonstrating CD33 inhibition of CD45. Together, these findings support that the CD33-CD45 interaction contributes to the pathology of AD.

**Figure 6.**
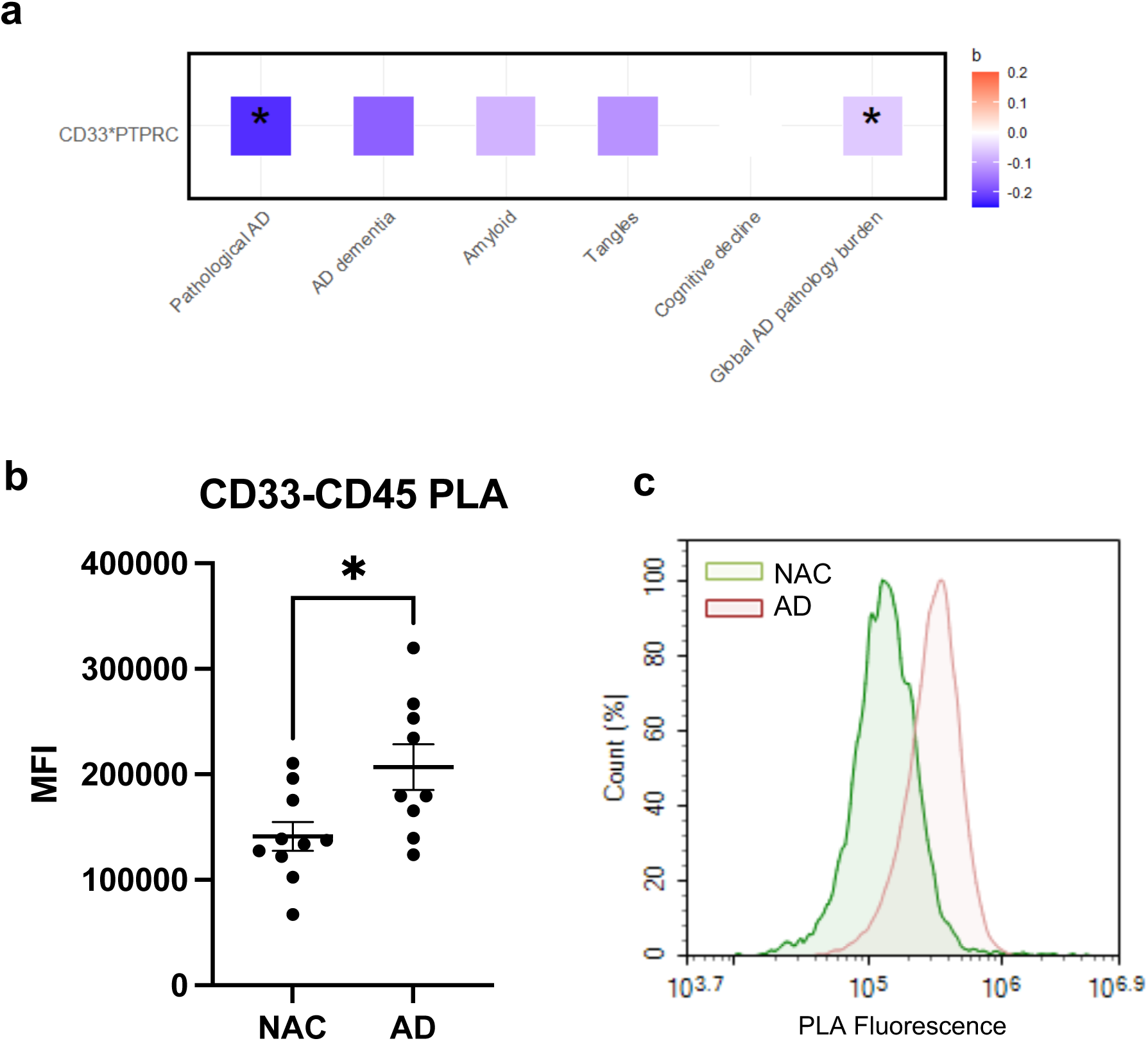
CD33-CD45 interaction frequency increases in Alzheimer’s disease, and CD33-PTPRC (CD45) gene expression interaction is associated with Alzheimer’s disease pathology. **a.** We investigated whether the interaction between *CD33* and *PTPRC* gene expression influences AD clinical and pathological traits using the transcriptomic dataset of dorsolateral prefrontal cortex from the ROSMAP cohort. After adjustment for multiple hypothesis testing, CD33-PTPRC interaction was significantly associated with pathological diagnosis of AD and global AD pathology burden, both with negative beta values. **b.** Median fluorescence intensity of CD33-CD45 proximity ligation assay (PLA) in primary monocytes from individuals with Alzheimer’s disease (AD) or non-AD controls (NAC). Donors were matched for sex and *CD33 rs3865444* genotype. N = 10 NAC donors and 9 AD donors. *p<0.05, unpaired t-test. **c.** Representative overlay flow cytometry histogram of PLA fluorescent signal from one AD and one NAC donor.

To better understand the relevance of this interaction in myeloid cells in AD, we next investigated whether this interaction is altered in primary monocytes from individuals with AD, in a genotype-independent manner. We performed PLA investigating CD33 and CD45 interaction in peripheral monocytes from 10 individuals with AD and 10 non-AD controls, with both groups matched for sex and *CD33 rs3865444* genotype (Supplementary Data Table 5). Interestingly, we observed increased CD33-CD45 PLA signal in monocytes from individuals with AD compared to controls (Figure 6b, 6c). These findings suggest that CD33-CD45 interactions increase in AD, supporting systemic changes in myeloid function in this disease. One potential explanation underlying this increased interaction in AD could be due to the increased glycosylation observed in disease^40–42^. Together, these findings support a role for the CD33-CD45 interaction in AD.

## Discussion

Since the *CD33* locus was associated with AD^9, 10^, we and others have interrogated the functional consequences of the risk-associated SNP to identify pathways through which CD33 contributes to AD^43^. We previously found the *CD33* risk allele to reduce uptake of amyloid-β^11, 27^, a finding which we have also shown here in CD33 knock-out HMC3 cells, which had increased internalization of amyloid-β. Our group and others determined that the *CD33* risk allele leads to decreased splicing of exon 2, which encodes for the sialic acid binding domain^12, 14^. Therefore, individuals with the risk allele have greater protein expression of the inhibitory, sialic-acid binding CD33^M^ isoform on the surface of their myeloid cells^11, 27^. However, what proteins CD33 interacts with through its sialic acid binding domain, and thereby regulates, are completely unknown and are of great interest to understand how increased CD33^M^ contributes to AD risk.

Here, we identify novel sialic acid-dependent binding partners of CD33 in innate immune cells, including CD45. We demonstrate increased CD33-CD45 interaction in individuals homozygous for the *CD33* AD risk allele in peripheral monocytes and human brain tissue, which is presumably due to the genetically dictated increase of the sialic acid binding isoform. Functionally, we identified that CD33 suppresses CD45 phosphatase activity. Relevant to microglial function in AD, this interaction increased following Aβ1-42 exposure. CD33^M^ overexpression and CD45 inhibition impaired microglial clearance of Aβ_1-42_ and contributed to amyloid-induced synaptic loss, supportive of a detrimental effect of CD33-mediated inhibition of CD45 phosphatase activity in the context of AD pathology^28–30^. Interestingly, the CD33-CD45 interaction was increased in monocytes from individuals with AD compared to controls, in a genotype-independent manner, suggesting this interaction participates in systemic changes to myeloid function in this disease. Together, our findings suggest that CD33-mediated CD45 phosphatase inhibition may contribute to the AD risk associated with these molecules by impairing protective myeloid functions in disease.

Siglecs are sialic acid-binding receptors on the surface of immune cells, which are thought to regulate the innate immune inflammatory response. Siglecs are known to engage in *cis* interactions, which are thought to regulate inflammatory responses through inhibitory signal transduction^44, 45^. Indeed, desialylation has been shown to lead to increased inflammatory activity in THP-1 monocytes and human neutrophils^46^. However, the AD genetic risk associated with the sialic acid binding site of CD33 suggests that this inhibitory signaling may chronically contribute to disease processes. Our findings suggest that the inhibitory *cis* interaction between CD33 and CD45 in microglia may be maladaptive and impair microglial clearance of amyloid pathology. As CD33 may have an important role in regulating innate immune inflammatory responses, therapeutically blocking or downregulating CD33 may cause excess inflammation^47^. However, impairing CD33 binding to its signaling partners, through approaches such as competitive inhibition or neutralizing the receptor, may provide a practical approach to targeting CD33 interactions in disease.

CD33 is the only siglec with a SNP specifically associated with AD, suggesting that CD33 may modulate AD risk via a mechanism unique to CD33 or that CD33 may be the only siglec with a genetic variant that alters its function^6^. In keeping with this idea, we have previously demonstrated increased interaction between CD33 and SHP-1, CD33’s downstream phosphatase signaling partner, in individuals homozygous for the AD-risk variant^8^. Our present findings similarly demonstrate a genotype effect modulating the CD33-CD45 interaction, such that individuals homozygous for the AD-risk variant have increased interaction. While CD45 is also known to bind CD22, another siglec not expressed on human microglia, CD45’s interaction with other siglecs expressed on microglia has not been previously reported.

CD45 is a protein tyrosine phosphatase expressed on all immune cells, including the majority of human adult microglia^3, 48^, which dephosphorylates Src kinases^49^. The role of CD45 in B and T cell receptor (TCR) signaling has been well established, in which it has both negative and positive regulatory roles. In T cells, CD45 activates the Src family kinase Lck, which phosphorylates the TCR complex. CD45 has also been found to suppress TCR signaling in the context of low-affinity antigen or antigen-independent signaling by dephosphorylating signaling motifs within the TCR complex^50^. However, its role in innate immune cells is less known. Several CD45 substrates have been identified, including DAP12, Janus kinases, and Src family kinases^51^. In innate immune cells, CD45 has been shown to negatively regulate JAK/STAT signaling^52^ and participate in cell motility^53^. However, the specificity of this regulation remains unclear.

We report here that CD45 is a sialic acid-dependent binding partner of CD33, and that this interaction leads to a suppression of CD45 phosphatase function, in keeping with the known inhibitory function of Siglecs. We found that mutating CD33 at the ITIM motif abrogated the interaction between CD33 and CD45, suggesting that CD33-CD45 interaction requires two aspects: 1) extracellular binding at the sialic acid binding domain and 2) active CD33 signaling through the ITIM motif, via molecules such as SHP-1, which we have previously shown to interact with CD33 in a genotype dependent manner^8^. CD33 intracellular signaling may facilitate a complex of other partners, including SHP-1 and Src family members, which are known to interact with CD45. In this way, a cytoplasmic signaling cascade may be important to facilitate extracellular CD33-CD45 binding and subsequent CD45 phosphatase inhibition.

CD45/*PTPRC* was identified as a key driver of AD by transcriptomic analyses of human post-mortem brain tissue^20, 22^. Supportive of a detrimental role for reduced CD45 activity in the setting of AD pathology, when PS/APP (presenilin and APP) AD transgenic mice were crossed with CD45-/- mice, CD45 deficiency was found to accelerate cerebral amyloidosis and increase neuronal loss. The CD45-deficient murine microglia were also found to be defective in amyloid-β uptake^23^. These findings are consistent with our results demonstrating a loss of amyloid-β internalization in CD45 knock-out HMC3 cells, conferring vulnerability to amyloid-β-induced spine loss. Our findings further demonstrate that CD33^M^ overexpression mirrored the same impairment in phagocytosis and amyloid-β induced spine loss as CD45 knock-out, aligning with our findings of CD33 ^M^-mediated inhibition of CD45.

In AD, we found increased CD33-CD45 interactions on monocytes, independent of CD33 genotype, further suggesting a role for increased CD33 interaction with CD45 in systemic myeloid dysregulation in this disease. This increased interaction may be due to increased glycosylation that has been reported in AD^40–42^. Furthermore, this interaction increased following exposure to amyloid-β, suggesting that CD33-CD45 interaction, while increased in CD33 risk carriers, also increases in the context of AD pathology. Functionally, we demonstrated that CD33^M^ overexpression and CD45 inhibition, aligning with CD33^M^-mediated inhibition of CD45, impairs protective microglial functions, including maintenance of neuronal health and clearance of amyloid-β; impairments that may contribute to risk for disease.

The focus of this work was to identify binding partners of CD33 produced by innate immune cells. In our companion paper, Dodd *et al*. identified clusterin as a soluble sialic acid-dependent binding partner for CD33. Clusterin, also genetically associated with AD, is predominantly produced by astrocytes and therefore represents crosstalk between different cell types involving AD genetically implicated proteins^54^. Our findings suggest that clusterin enhances the interaction of CD33-CD45. Future work may investigate how trans-acting ligands (such as clusterin) and cis-acting ligands (such as CD45) interact to coordinately determine CD33 activation status, with the ultimate goal of elucidating mechanisms through which sialic acid-dependent CD33 activity confers risk for AD.

## Materials and Methods

### Cell culture (THP-1, monocytes, MDMi)

The THP-1 cell line was cultured in RPMI-1640 Glutamax (Life Technologies) medium containing 10% fetal bovine serum (FBS; Corning), 20 mM HEPES (Lonza), 1mM Sodium Pyruvate (Lonza), 1X MEM-NEAA (Life Technologies), and supplemented with 1% Penicillin-Streptomycin (Life Technologies).

PBMCs from healthy individuals from the PhenoGenetic Project or the New York Blood Center were selected based on their genotype for *rs3865444*. Informed consent was obtained from all human subjects. All blood draws and data analyses were done in compliance with protocols approved by the institutional review boards of each institution. PBMCs were separated by Lymphoprep gradient centrifugation (StemCell Technologies). PBMCs were frozen at a concentration of 1x10^7^ cells/ml in 10% dimethyl sulfoxide (DMSO; Sigma Aldrich) and 90% (v/v) fetal bovine serum (FBS; Corning). Before each study, aliquots of frozen PBMCs were thawed and washed in 10 ml of phosphate-buffered saline (PBS). Monocytes were positively collected from whole PBMCs using anti-CD14+ microbeads (Miltenyi Biotec) and plated at 1.5 x 10^5^ cells per well in 96-well plates. To induce the differentiation of MDMi, we incubated monocytes under serum-free conditions using RPMI-1640 Glutamax (Life Technologies) with 1% Penicillin-Streptomycin (Life Technologies) and Fungizone (2.5 μg/ml; Life Technologies) and a mixture of the following human recombinant cytokines: M-CSF (10 ng/ml; Biolegend, 574806), GM-CSF (10 ng/ml, R&D Systems, 215-GM-010/CF), NGF-β (10 ng/ml, R&D Systems, 256-GF-100), CCL2 (100 ng/ml; Biolegend, 571404), and IL-34 (100 ng/ml; R&D Systems, 5265-IL-010/CF) under standard humidified culture conditions (37°C, 5% CO_2_) for 10 days^26^.

### Photocrosslinking

Ac_4_ManNDAz was prepared as described previously^17, 18^. THP-1 cells were seeded at 0.4 x 10^6^ cells/mL and cultured with DMSO or 0.1 mM Ac_4_ManNDAz for 3 days (0.1% DMSO final concentration), harvested, and resuspended in ice cold PBS at 2.5 x 10^6^ cells/mL. They were then irradiated for 10 mins at 365 nm on ice in clear 6-well plates (Corning) with a UV irradiator. Cells were then pelleted and lysed in RIPA buffer with protease and phosphatase inhibitors.

### Immunoprecipitation (IP) and Mass Spectrometry (MS)

For MS experiments, 50 x 10^6^ cells were harvested after Ac_4_ManNDAz or DMSO treatment, irradiated or left untreated, and lysed as above. For immunoprecipitation MS experiments, lysates (5 mg) were incubated with 25μg of anti-CD33 antibody (WM-53, Biolegend) overnight with end-over-end rotation at 4°C. 250 μL of Protein G Dynabeads (Invitrogen) were resuspended, the supernatant was removed, and then the above lysate and antibody was added. After 8 hours with end-over-end rotation at 4°C, beads were washed three times with TBS, resuspended in 10μL of TBS, and immediately frozen at - 80°C until MS processing.

#### Protein digestion and labeling with TMT isobaric mass tags

The beads from immunopurification were washed once with IP lysis buffer, then three times with PBS. The lysates of each replicate were resuspended in 90μL digestion buffer (2M Urea, 50 mM Tris HCl) and 2μg of sequencing grade trypsin was added, with 1 hour shaking at 700rpm. The supernatant was removed and placed in a fresh tube. The beads were then washed twice with 50μL digestion buffer and combined with the supernatant. The combined supernatants were reduced (2μL 500 mM Dithiothreitol (DTT), 30 minutes, RT), alkylated (4μL 500 mM Iodoacetamide (IAA), 45 minutes, dark) and a longer overnight digestion performed: 2μg (4μL) trypsin, with shaking overnight. The samples were then quenched with 20μL 10% formic acid (FA) and desalted on 10mg Oasis cartridges. Desalted peptides were labelled with TMT10 reagents (Thermo Fisher Scientific, lot QL228730) according to the table below. Peptides were dissolved in 25μL of fresh 100mM HEPES buffer. The labelling reagent was resuspended in 42μL of acetonitrile and 10μL added to each sample as described below. After 1 h incubation the reaction was stopped with 8μL of 5mM Hydroxylamine.

Labelling Scheme:

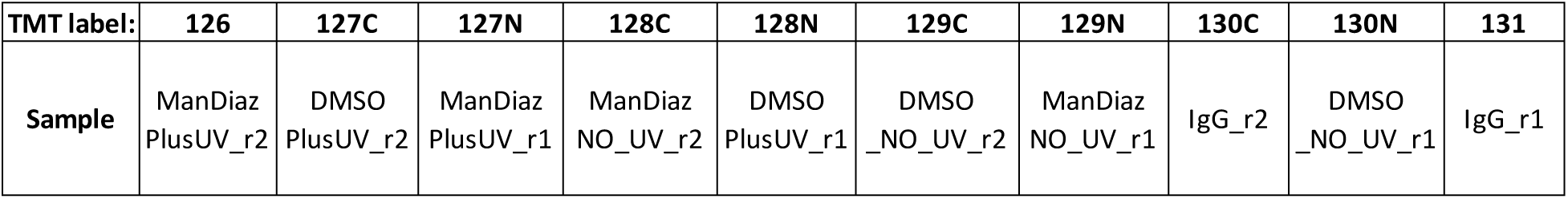

#### Protein identification with nanoLC–MS system

Reconstituted peptides were separated on an online nanoflow EASY-nLC 1000 UHPLC system (Thermo Fisher Scientific) and analyzed on a benchtop Orbitrap Q Exactive plus mass spectrometer (Thermo Fisher Scientific). The peptide samples were injected onto a capillary column (Picofrit with 10 μm tip opening / 75 μm diameter, New Objective, PF360-75-10-N-5) packed in-house with 20 cm C18 silica material (1.9 μm ReproSil-Pur C18-AQ medium, Dr. Maisch GmbH) and heated to 50°C in column heater sleeves (Phoenix-ST) to reduce backpressure during UHPLC separation. Injected peptides were separated at a flow rate of 200 nL/min with a linear 230 min gradient from 100% solvent A (3% acetonitrile, 0.1% formic acid) to 30% solvent B (90% acetonitrile, 0.1% formic acid), followed by a linear 9 min gradient from 30% solvent B to 60% solvent B and a 1 min ramp to 90%B. Each sample was run for 260 min, including sample loading and column equilibration times. The Q Exactive instrument was operated in the data-dependent mode acquiring HCD MS/MS scans (R=17,500) after each MS1 scan (R=70,000) on the 12 topmost abundant ions using an MS1 ion target of 3x10^6^ ions and an MS2 target of 5x10^4^ ions. The maximum ion time utilized for the MS/MS scans was 120 ms; the HCD-normalized collision energy was set to 27; the dynamic exclusion time was set to 20s, and the peptide match and isotope exclusion functions were enabled.

#### Database search and data processing

All mass spectra were processed using the Spectrum Mill software package v6.0 pre-release (Agilent Technologies) which includes modules developed by the Carr laboratory for TMT10-based quantification. For peptide identification, MS/MS spectra were searched against human Uniprot database to which a set of common laboratory contaminant proteins was appended. Search parameters included: ESI-QEXACTIVE-HCD scoring parameters, trypsin enzyme specificity with a maximum of two missed cleavages, 40% minimum matched peak intensity, +/- 20 ppm precursor mass tolerance, +/- 20 ppm product mass tolerance, and carbamidomethylation of cysteines and TMT6 labeling of lysines and peptide n-termini as fixed modifications. Allowed variable modifications were oxidation of methionine, N-terminal acetylation, Pyroglutamic acid (N-termQ), Deamidated (N),Pyro Carbamidomethyl Cys (N-termC), with a precursor MH+ shift range of -18 to 64 Da. Identities interpreted for individual spectra were automatically designated as valid by optimizing score and delta rank1-rank2 score thresholds separately for each precursor charge state in each LC-MS/MS while allowing a maximum target-decoy-based false-discovery rate (FDR) of 1.0% at the spectrum level.

TMT10 reporter ion intensities were corrected for isotopic impurities in the Spectrum Mill protein/peptide summary module using the afRICA correction method which implements determinant calculations according to Cramer’s Rule^55^ and correction factors obtained from the reagent manufacturer’s certificate of analysis (https://www.thermofisher.com/order/catalog/product/90406) for lot number QL228730.

### Proximity Ligation Assay (PLA)

The PLA assay was performed as per the manufacturer’s instructions (Sigma Aldrich). Briefly, THP-1 cells or MDMi were fixed in 4% Paraformaldehyde (PFA) (Electron Microscopy Sciences) for 15 minutes at room temperature. They were then permeabilized in PBS containing 1% Triton X-100 (Sigma Aldrich) for 15 minutes before being blocked by 10% normal goat serum in PBS with 1% Triton for 30 minutes. In the first antibody incubation, rabbit anti-human CD33 (Sigma Aldrich, HPA035832, 1:400) and mouse anti-human CD45 (Novus, NB500-319, 1: 400) were performed overnight at 4°C. After two washes with buffer A (Sigma Aldrich), secondary antibodies conjugated with oligonucleotides (PLA probe MINUS and PLA probe PLUS, Sigma Aldrich) were added to the reaction and incubated for one hour. The samples were then washed with Buffer A two times. The Ligation solution, consisting of two oligonucleotides and Ligase, was added and the oligonucleotides hybridized to the two PLA probes. Then the samples were washed with Buffer A two more times. The Amplification solution, consisting of nucleotides and fluorescently-labeled oligonucleotides, were added together with the polymerase. The oligonucleotide arm of one of the PLA probes acts as a primer for a rolling-circle amplification (RCA) reaction using the ligated circle as a template, generating a concatemeric (repeated sequence) product. The fluorescently-labeled oligonucleotides will hybridize to the RCA product. The samples were washed with Buffer B and then mounted with Duolink In Situ Mounting Medium with DAPI (Sigma Aldrich). The signal is visible as a distinct fluorescent spot and analyzed by confocal (Zeiss LSM 700) and fluorescence (Zeiss Axio) microscopies. PLA Images were analyzed in ImageJ. PLA immunofluorescent signal was thresholded and quantified by area, then normalized by the number of DAPI+ cells.

### Flow PLA

The Flow-PLA assay was performed as per the manufacturer’s instruction (Sigma Aldrich). Briefly, primary monocytes were fixed for 30 minutes on ice using the Invitrogen Fix & Perm Cell Permeabilization Kit. In the first antibody incubation, rabbit anti-human CD33 (Sigma Aldrich, HPA035832, 1:400) and mouse anti-human CD45 (Novus, NB500-319, 1: 400) were performed overnight at 4°C. After two washes with Duolink wash buffer (Sigma Aldrich), secondary antibodies conjugated with oligonucleotides (PLA probe MINUS and PLA probe PLUS, Sigma Aldrich) were added to the reaction and incubated for one hour at 37°C. The samples were then washed with Duolink wash buffer twice. The Ligation solution, consisting of two oligonucleotides and Ligase, was added and the oligonucleotides hybridized to the two PLA probes. Then the samples were washed with Duolink wash buffer two more times. The Amplification solution, consisting of nucleotides and fluorescently-labeled oligonucleotides, were added together with the polymerase. The oligonucleotide arm of one of the PLA probes acts as a primer for a rolling-circle amplification (RCA) reaction using the ligated circle as a template, generating a concatemeric (repeated sequence) product. The fluorescently-labeled oligonucleotides will hybridize to the RCA product. The samples were washed with Duolink wash buffer and then incubated with the Duolink flowPLA Violet Detection solution. The signal was analyzed by flow cytometry (Novocyte Quanteon Cell Analyzer).

### In Situ PLA

Autopsy tissue was obtained from the New York Brain Bank at Columbia University (Supplementary Data Table 3). Samples were collected in compliance with protocols approved by the Columbia Human Research Committee. The genotype for CD33 was determined using Custom Taqman SNP Genotyping Assay (ThermoFisher Scientific, Assay ID AHLJYS8) on the QuantStudio 7 Flex Real-Time PCR System. The frozen tissues were thawed at room temperature for 10 min after being taken from -80°C. The tissues were fixed with pre-cooled 100% ethanol at -20°C for 15 min. After being washed three times with PBS, the tissues were immersed in PBS for 10 min and then blocked in 3% Bovine serum albumin (BSA) with 0.1% Triton-X for 1 hr. Tissues were washed three times for 5 min each in PBS. Incubation with primary antibodies was performed overnight at 4°C in 1% BSA. The PLA secondary probes were diluted 1:5 in antibody diluent and incubated for 20 min at room temperature. The samples were washed three times with PBS. The slides were incubated in the probe mixture at 37°C for 1.5 hours in a humidified chamber, followed by washing with DuoLink buffer A two times and incubation with ligation solution at 37°C for 1.5 hours in a humidified chamber and washed with Buffer A two times for 10 min at room temperature. The slides were then incubated with amplification-polymerase solution at 37°C for 100 min in a humidified chamber and washed with Buffer B two times for 10 min at room temperature. To identify microglia, slides were incubated with anti-CD45 (Novus, MEM-28) at a 1:100 dilution for 1.5 hours and washed three times in PBS followed by an incubation with the secondary antibody for 1 hour and washed three times with PBS. Slides were incubated with Sudan Black B (1% w/v in 100% ethanol, Sigma Aldrich) for 3 min to quench the autofluorescence and washed three times with PBS. Finally, the sections were mounted with Duolink In Situ Mounting Medium with DAPI and imaged with an immunofluorescence microscope.

### Flow Cytometry Protein Staining

Cells (THP1, primary monocytes, MDMi, or HMC3) were stained on ice with CD33-APC (Miltenyi, cat. 130-113-345, 1:50) or CD45-PE (BioLegend, cat. 304008, 1:20) for 30 minutes away from light according to the manufacturer. Isotype controls (CD33: Myltenti, cat. 130-113-196l; CD45: BioLegend, cat. 400112) were similarly stained as controls. Cells were then washed three times with PBS with 1% fetal calf serum and labeled with Fixable LIVE/DEAD violet cell stain (Thermo Fisher Scientific) for 30 minutes and fixed in 4% paraformaldehyde (vol/vol) for 30 min. For HMC3 cultures, cells were treated with trypsin 48 hours prior to staining and plated in a polypropylene plate in which staining was later performed. For extended data figure 9, HMC3 cells were incubated with the eBioscience Intracellular Fixation and Permeabilization kit (Invitrogen) for 30 minutes and then washed prior to staining. All flow cytometry data were collected on a NovoCyte Quanteon cell analyzer and compensated and analyzed using the associated NovoExpress software.

### Knockout Cell Generation

THP-1-Cas9 or HMC3-Cas9 expressing cells were generated via lentiviral integration and blasticidin selection of single cell clones using the Lenti-Cas9-Blast plasmid (Addgene 52962). gRNAs for CD33 and CD45 were designed using the Broad GPP webportal and cloned into the pLKO-sgRNA-EFS-GFP vector (52962) using established protocols^56^. Non-integrating lentiviruses were then generated using Lenti-X Packaging Single Shots (Clontech) and used to infect the above cells. Infection and deletion were assessed using GFP and cell-surface staining. Knockout cells thereby generated were then sorted by flow cytometry and single cell clones were generated by limiting dilution.

gRNA:

CD33 CRISPR F1 Sequence: GCCACTCACCTGCCCACAGC CD33 CRISPR R1 Sequence: GGCTACTGCTGCCCCTGCTG CD45 CRISPR F1: CACCGGAAACTTGCTGAACACCCG CD45 CRISPR R1: AAACCGGGTGTTCAGCAAGTTTCC

### CD33 Mutant Cell Lines

A CD33 encoding lentivirus was obtained from Origene (RC207023L1) and various CD33 mutants were made (R119K, Y340F, Y358F, and Y340F/Y358F) using the Q5 Site-Directed Mutagenesis Kit (New England Biolabs). Lentiviruses were then transfected into CD33 KO cells and cells with similar expression levels were then sorted using flow cytometry.

Primers:

R119K F Primer: ATACTTCTTTAAGATGGAGAGAGGAAG

R119K Reverse: GAACCATTATCCCTCCTC

Y340F F: GGAGCTGCATTTTGCTTCCCTCA

Y340F R: TCATCCATCTCCACAGTAGG

Y358F F: CTCCACCGAATTCTCAGAGGTCAG

Y358F R: GTGTCCTTGGAAGGATTCATC

### Dot blot assay

Protein samples were diluted in Dilution buffer (8M Urea, 100mM NaH_2_PO_4_, 10mM TrisHCl, pH 8.0) to final protein concentrations of 100ng/ml). A circle on the Nitrocellulose membrane (Biorad) was drawn with a pencil. 1ml samples of diluted protein were loaded directly onto the membrane inside the circle. The membrane was given time to dry for a short time at room temperature. The membrane was next blocked in Blocking buffer (1% BSA in PBST) for 1 hour at room temperature on a shaker and then washed three times (10 minutes per wash) with Wash buffer (PBST:0.1% Tween-20 in PBS). The primary antibody (CD45 antibody (MEM-28), Novus Biologicals, NB500-319) was diluted in Blocking buffer at 1:5000. The membrane was incubated in 10ml pre-diluted primary antibody for 1 hour at room temperature on a gentle shaker. The membrane was washed three times (10 minutes per wash on a gentle shaker) with Wash buffer (PBST:0.1% Tween-20 in PBS). The secondary antibody (goat anti-mouse IgG (H+L) HRP, Invitrogen, 31431) was diluted in Blocking buffer at 1:10000. The membrane was washed three times (10 minutes per wash on a gentle shaker) with Wash buffer (PBST:0.1% Tween-20 in PBS). The membrane was then placed on a plastic board, covered with Saran wrap and incubated with 1000 ml ECL substrate reagent (Pierce™ ECL Western Blotting Substrate, Thermo Fisher, 32106) for 5 minutes. The substrate was discarded, and the remaining liquid was soaked up with soft paper. The membrane was visualized for 30 seconds using the BIO-RAD® ChemiDocTM MP Imaging System. The white-light picture was used to double-check the results. The same procedure was repeated in the same membrane for loading control with primary antibody (beta Actin Polyclonal antibody, Invitrogen, PA1-183) and secondary antibody (Goat anti-rabbit IgG (H+L), HRP, Invitrogen, 31460)

### Agarose gel electrophoresis of PCR products

Genomic DNA (gDNA) was extracted using Blood & Cell Culture DNA Mini Kit (Quiagen, 13323) according to the manufacturer’s protocol. Concentration of gDNA was assessed using the NanoDrop spectrophotometer (Thermo Scientific, 701-059049) following manufacturer’s instructions. To generate PCR amplicons across the target site, *CD33* exon 1 locus-specific primers, forward 5’-CTCCTCCCCAGCTTCCTGTC-3’ and reverse 5’- GTGGGGAAACGAGGGTCAGCT-3’ were designed. The amplicon size was 650 bp around the predicted Cas9 cutting site. For PCR, 30–40 ng of gDNA input were used as a template for DreamTaq PCR Master Mixes (2X) (Thermo Fisher, K1081) with the following thermocycler conditions: denaturing for 3 minutes at 95°C; then 35 cycles of 1 minute at 95°C, primer annealing for 30 seconds at 60°C and amplification of 1 minute at 72°C, and a final extension for 5 minutes at 72°C. Amplicons were analyzed by 1.5% TBE agarose gel electrophoresis.

### CD33-CD45 Solid Phase Binding Assay

To measure the binding of CD33 to CD45, a solid-phase-based binding assay was performed as previously described^57^. Briefly, recombinant human CD45 (R&D Biosystems, 1430-CD-050) was diluted in carbonate-bicarbonate buffer (pH=9.6) to concentrations of 0, 0.5μg/mL, or 1μg/mL and 50μL/ well was added to a 96 well plate and incubated overnight at 4°C to immobilize CD45 to the plate. Wells were then blocked with 100μL of 1% bovine serum albumin for 1 hour at room temperature and washed 3 times with 200μL per well of HEPES buffered saline + 1% Tween-20 (HBS-T). 100μL of increasing concentrations of recombinant hCD33-Fc (R&D Biosystems, 1137-SL-050) was then incubated on the plate overnight at 4°C. Recombinant hCD33-Fc consists of the extracellular domain of human CD33 containing the sialic acid binding domain fused to the Fc domain of human IgG. Following wash steps with HBS-T to reduce non-specific binding, plates were then incubated with 100mL of anti-hIgG Fc-HRP conjugated antibody (Thermo Fisher, A18817) at a 1:10000 dilution for 1 hour at room temperature to detect bound CD33. Following 3 washes with HBS-T, plates were then incubated with Quantablu fluorescent substrate for 30 minutes before reading fluorescence at 320nm/450nm on a Tecan Infinite 200 Pro microplate reader. A binding curve was established by varying the concentrations of hCD33-Fc in the assay, and the resulting changes in fluorescent intensity were measured as an indicator of bound hCD33-Fc.

To determine sialic acid dependent binding, prior to incubating wells with recombinant hCD33, 20 μL of sialidase (neuraminidase, Sigma-Aldrich, 11080725001) at 500mU/mL was added to each well and incubated at 37°C for 1 hour. Wells were then washed 3x with HBS-T prior to adding hCD33.

### Phosphatase Activity Assay

5 x 10^5^ THP-1, THP-1 CD33KO, THP-1 CD33R119K, or THP-1 CD33WT cells were pre-incubated with or without LPS (10 ng/ml) (Invitrogen) at 37°C for 18 hours and CD45 phosphatase activity was determined as described, with modifications^57^. Briefly, cells were washed with ice-cold PBS and lysed in Ph lysis buffer (20nM HEPES (Fisher scientific), pH 7.2, 2 mM EDTA (Invitrogen), 2mM dithiothreitol (Sigma Aldrich), 1% Nonidet P40 Substitute solution (Sigma Aldrich) and 10% (v/v) Glycerol (Sigma Aldrich) containing complete, Mini, EDTA-free Protease Inhibitor Cocktail (Roche). Cellular debris and nuclear material were removed by centrifugation at 20,000 x g for 15 min at 4°C. Phosphatase activity was determined by incubation of 20 μg of lysed proteins for 4 hours at 37°C with 1 mg/ml 4-Nitrophenyl phosphate disodium salt hexahydrate (Sigma Aldrich) in CD45 Ph assay buffer (100 nM HEPES, pH 7.2, 2 mM EDTA and 2 mM dithiothreitol). The resulting color change was assessed at 410 nm.

### CD45 Specific Phosphatase Activity

5 x 10^5^ THP-1 cells were pre-incubated with or without anti-CD33 antibody WM53 (10 μg/mL) (Invitrogen, 14-0338-82) or IgG control (Biolegend, 400165) at 37°C for 48h. CD45 phosphatase activity was determined as described, with modifications^57^. Briefly, cells were washed with ice-cold PBS and lysed in Ph lysis buffer (20nM HEPES (Fisher scientific), pH 7.2, 2mM EDTA (Invitrogen), 2mM dithiothreitol (Sigma Aldrich), 1% Nonidet P40 Substitute solution (Sigma Aldrich) and 10% (v/v) Glycerol (Sigma Aldrich) containing complete, Mini, EDTA-free Protease Inhibitor Cocktail (Roche). Cellular debris and nuclear material were removed by centrifugation at 20,000 x g for 15 min at 4°C. Phosphatase activity was determined by incubation of 20μg of lysed proteins for 4 hours at 37°C with 1 mg/ml 4-Nitrophenyl phosphate disodium salt hexahydrate (Sigma Aldrich) in CD45 Ph assay buffer (100nM HEPES, pH 7.2, 2mM EDTA and 2 mM dithiothreitol). The resulting color change was assessed at 410 nm. Non-CD45 specific phosphatase activity was determined by the addition of CD45 Inhibitor VI (Sigma Aldrich, 5.30197) at 3μM for 3 hours prior to cell lysis. Phosphatase activity from cells treated with CD45 Inhibitor VI was subtracted from cells not treated with inhibitor to determine CD45 specific phosphatase activity.

### Clusterin Treatment

0.1mg of clusterin (BioVendor, Clusterin Human Native Protein, Human Plasma; Cat. No. 50-206-9196) was resuspended in 108μl of sterile water to make a 12μM stock. 1ul of the 12uM clusterin stock was added into 200μl of media per well in a 96-well plate for a final concentration of 60nM. The cells were then incubated with clusterin treatment for 15 minutes at 37°C. Cells were washed with PBS two times before performing CD33-CD45 PLA, as described above.

### Amyloid-beta Treatment

The Aβ_1-42_ was reconstituted in 1% of NH_4_OH in MilliQ water for 10 min at RT. Then, the solution was incubated at 37°C for 1 week with rotation. Cells at a concentration of 1x10^6^ were incubated with 250nM aggregated amyloid beta in RPMI medium for 24 hours at 37°C. Then the cells were washed with PBS two times before analysis by PLA, as described above.

### Primary Cortical Culture

Cortices from E15.5 mouse embryos, from timed pregnant CD1 mice (Crl:CD1(ICR)) obtained from Charles River, were dissected and electroporated, followed by dissociation in complete Hank’s balanced salt solution (cHBSS) containing papain (Worthington) and DNase I (100μg/mL, Sigma) for 15 minutes at 37°C, washed three times, and manually triturated in DNase I (100μg/mL) containing neurobasal medium (Life Technology) supplemented with B27 (1x, Thermo Fischer Scientific), FBS (2.5% Gibson), N2 (1x, Thermo Fischer Scientific), and GlutaMax (2mM, Gibco). Cells were plated at 1.0 x 10^5^ cells per 35mm glass bottom dish (MatTek) that had been coated with poly-D-lysine (1mg/mL, Sigma) overnight. One-half of the medium was changed every 5-7 days thereafter with non-FBS containing medium and maintained for 20-22 days at 37°C in 5% CO_2_.

### *Ex utero* Electroporation

*Ex utero* electroporation of mouse dorsal telencephalic progenitors was performed as previously described^36^ by injecting plasmid DNA (1-2 μg/μL endotoxin-free plasmid DNA, Midi prep kit from Macherey-Nagel) plus 0.5% Fast Green (1:20 ratio, Sigma) using the picospritzer III (Parker) microinjector into the lateral ventricles of isolated E15.5 embryonic heads that were decapitated and placed in complete HBSS^58^. Electroporation was performed on the whole head (skin and skull intact) with gold-coated electrodes (GenePads 5 × 7 mm BTX) using an ECM 830 electroporator (BTX) and the following parameters: five 100 ms long pulses separated by 100 ms long intervals at 20 V. Immediately after electroporation, cortices were prepared for primary neuronal culture.

### Primary Cortical and Human Microglia cell line Co-Culture

HMC3 cells were grown at 37°C in 5% CO_2_ in complete MEM containing FBS (10%, Gibson) with Pen-Strep (2%, Gibco) (and neomycin solution (2%, Santa Cruz) for the CD33^M^ line) were dissociated using Trypsin-EDTA (0.05%, Gibco), counted, and plated drop-wise onto 19DIV electroporated primary cortical mouse neurons, as described above, at 2.0 x 10^5^ cells per 35mm glass bottom dish (MatTek).

### Aβ1-42 Stock Preparation

HiLyteᵀᴹ Fluor 647 - labeled Beta - Amyloid1-42 (AnaSpec) was reconstituted in 50μL Ammonia Hydroxide and 3950 μl RPMI-1640 Glutamax (Life Technologies) medium. The stock was stored at -80 °C.

### Imaging

Imaging of dissociated neurons was performed between 20-22 DIV in 1,024 x 1,024 resolution with a Nikon Ti-E microscope equipped with A1R laser-scanning confocal microscope using the Nikon software NIS-elements (Nikon, Melville, NY, USA). We used the following objectives (Nikon): 40x (for analysis of microglial consumption) and 60x Apo TIRF; NA 1.49 (for analysis of spine densities of cultured neurons).

### Analysis of Spine Density

Dendritic spine densities were estimated on secondary or tertiary dendritic branches for cultured neurons using Fiji software (ImageJ, NIH). Spines were quantified over an average of 40μm in cultures, between 2-3 segments per cell. Spine density was defined as the number of quantified spines divided by the length over which the spines were quantified.

### Analysis of Microglial Consumption

Using Nikon software NIS-elements (Nikon, Melville, NY, USA), a binary threshold for each individual channel (FITC = Glia, Cy5 = Aβ) was set for each z-slice imaged. Upon binarization, the area occupied by signal was quantified in μm^2^ for each channel individually for each z-slice. Subsequently, the area of any co-localization between the FITC channel and Cy5 channel for each z-slice was quantified in μm^2^. In doing so, a 3D volumetric representation of each microglia was captured as a series of 2D area slices, in addition to a 3D volumetric representation of the Aβ. The total volume of microglial space, represented by stacked 2D area, occupied by Aβ (Cy5) was then calculated by taking the 2D area of co-localization—i.e. the area of space occupied by all Aβ signal within a single z-slice—and dividing it by the total possible 2D area—ie the area of space occupied by all microglia within a single z-slice—of the microglial population, and repeating for each z-slice for the image. This produces a curve representative of the area of 3D microglial space occupied by Aβ, specifically measuring intracellular Aβ.

### Gene-gene interaction analysis

To explore the interplay between CD33 and CD45 genes, we utilized previously described RNA-sequencing data from the DLPFC in ROSMAP study^59, 60^. All ROSMAP participants were enrolled without known dementia and agreed to detailed clinical evaluation and brain donation at death^39^. Clinical and pathologic methods have been previously described^61–66^.

Linear regression models for quantitative traits and logistic regression models for binary traits were used to test the interaction between CD33 and CD45 gene expression on (a) neuropathological AD status (pathologic AD), (b) β-amyloid load, (c) tau tangle density (tangles), (d) global measure of pathology based on the scaled scores of 5 brain regions (Global AD pathology burden), (e) estimated slope of global cognition using longitudinal measurements (cognitive decline), and (f) clinical dementia diagnosis antemortem (AD dementia). The gene expression levels were adjusted for age, sex, RIN score, post-mortem interval, and other technical covariates for RNA-sequencing and the residual expression values were used for interaction testing.

## Acknowledgements

The authors are grateful to the participants of the NYBC and NYBB for the time and specimens that they contributed. We would like to thank Franck Polleux for his support and advice. Research reported in this publication was performed in the Columbia Center for Translational Immunology (CCTI) Flow Cytometry Core, supported in part by the Office of the Director, National Institutes of Health under award S10RR027050. We thank the Columbia University Alzheimer’s Disease Research Center, funded by NIH grant P30AG066462 to S.A. Small (P.I.), and A. Teich for providing biological samples and associated information. We thank the National Centralized Repository for Alzheimer’s Disease and Related Dementias (NCRAD), supported by the National Institutes of Health under award U24AG021886. This work was supported by the US National Institutes of Health grants RF1AG058852, R01AG043617, R01GM090271, U01AG046152, R01AG048015, F30AG074618, F31AG079613, and T32AI148099. ROSMAP is supported by P30AG10161, P30AG72975, R01AG15819, R01AG17917, U01AG46152, and U01AG61356. The content is solely the responsibility of the authors and does not necessarily represent the official views of the National Institutes of Health. AJL, BNV, and EMB are all current Ludwig Scholars in the Carol and Gene Ludwig Center for Research in Neurodegeneration.

## Author Contributions

N.V., C.D.R., D.M.V. Z.K.C., and E.M.B. implemented the study and wrote the manuscript, and all authors read and edited the manuscript. Mass spectrometry experiments were designed and executed by C.D.R., P.L.D., G.G., J.J.K., M.S., S.A.C, and E.M.B. N.V., M.T., S.M.C., M.R., S.C., Z.K.C., J.L.H, K.A.T., P.S.G.H, and E.M.B designed and conducted the PLA experiments. N.V., J.L.H., W.E., and E.M.B., designed and conducted the phosphatase experiments. D.M.V., N.V., W.E., and E.M.B. designed and conducted the co-culture experiments. M.L., A.J.L., B.N.V., and D.A.B were responsible for the gene-gene interaction experiments in the ROSMAP dataset.

## Citation for the Submitted manuscript

The original mass spectra will be deposited in the public proteomics repository MassIVE and will be accessible at ftp://MSV000012345@massive.ucsd.edu when providing a dataset password and username. This data will be made public upon acceptance of the manuscript.

