## Extended Data Figures for "CD33-CD45 Interaction Reveals a Mechanistic Link to Alzheimer’s Disease Susceptibility"

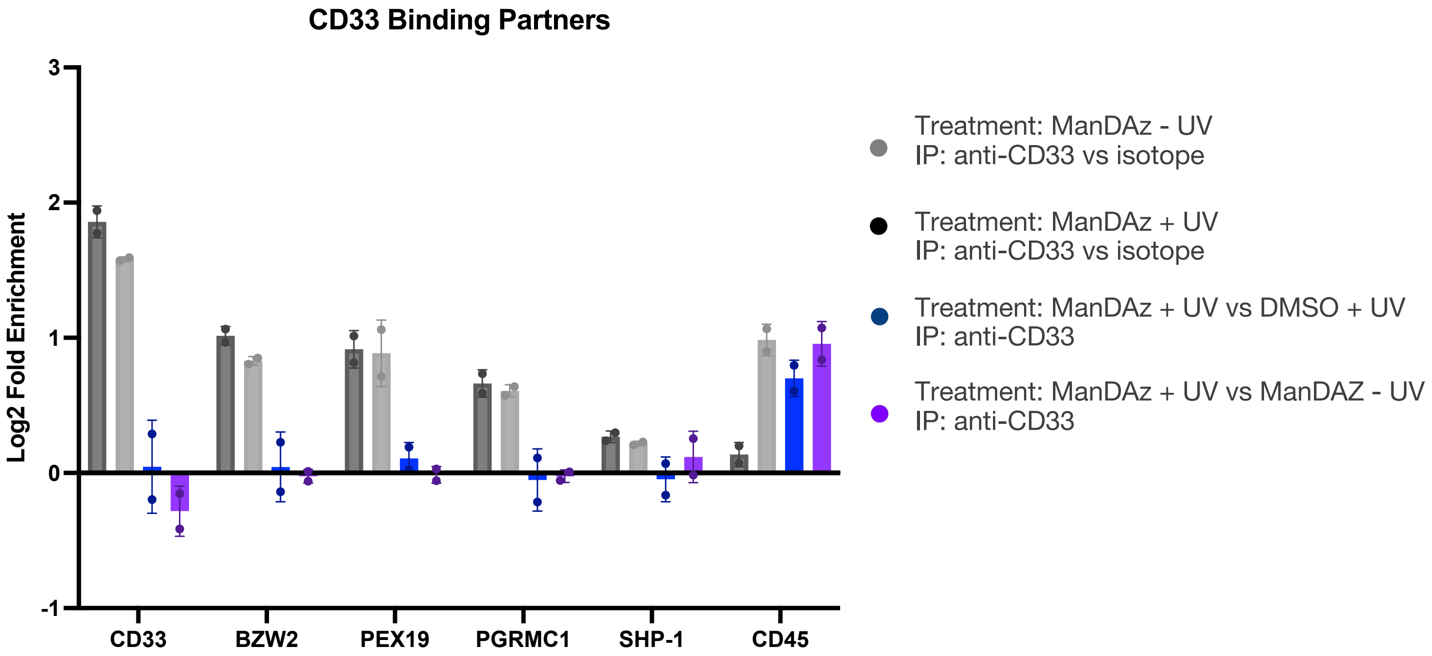


**Extended Data Figure 1: NanoLC-MS identifies binding partners of CD33 that are not sialic acid specific.** These proteins were enriched in the CD33 immunoprecipitation with Ac4ManNDAz with and without UV crosslinking compared to IgG control immunoprecipitation, demonstrating they do bind specifically to CD33. However, they do not interact specifically through the sialic acid binding domain as they are not enriched in the Ac4ManNDAz plus UV condition compared to either DMSO alone plus UV or Ac4ManNDAz without UV crosslinking. Error bars denote the standard deviation between technical replicates. CD45 is included as an example of a sialic acid dependent binding partner.

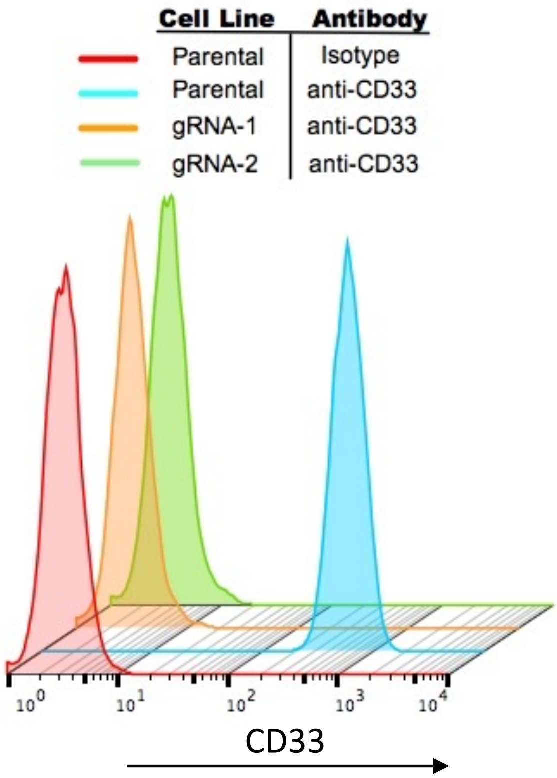


**Extended Data Figure 2. CD33 expression in WT and CD33 knock out (KO) THP-1 monocytes.** In CD33KOs, CD33 was deleted from the THP-1 cell line using Crispr/Cas9, and reduction in CD33 protein expression was validated by flow cytometry.

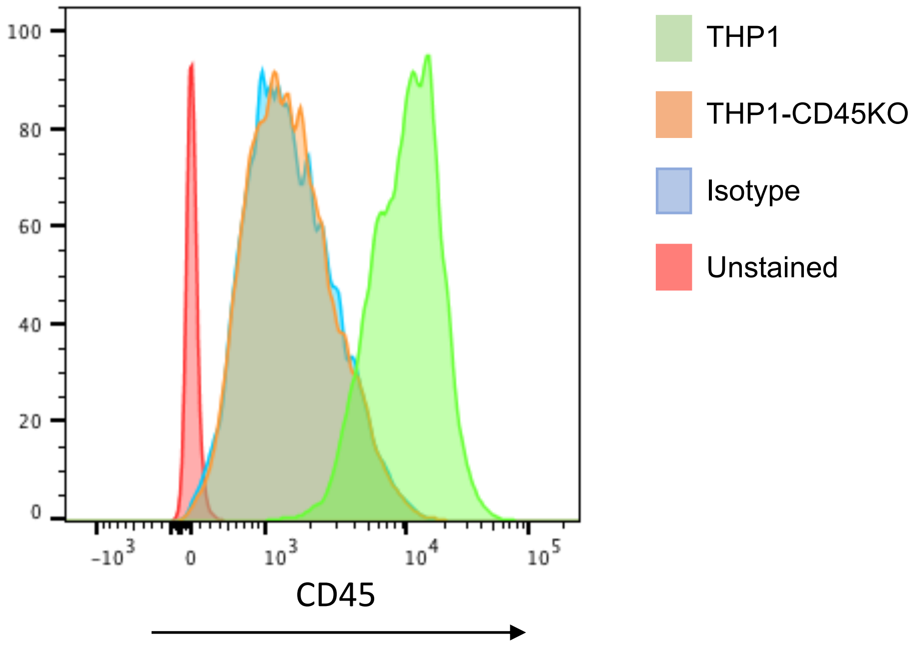


**Extended Data Figure 3. CD45 expression in WT and CD45 knock-out (KO) THP-1 monocytes.** In CD45KOs, expression was deleted from the THP-1 cell line using Crispr/Cas9, and reduction in CD45 protein expression was validated by flow cytometry.

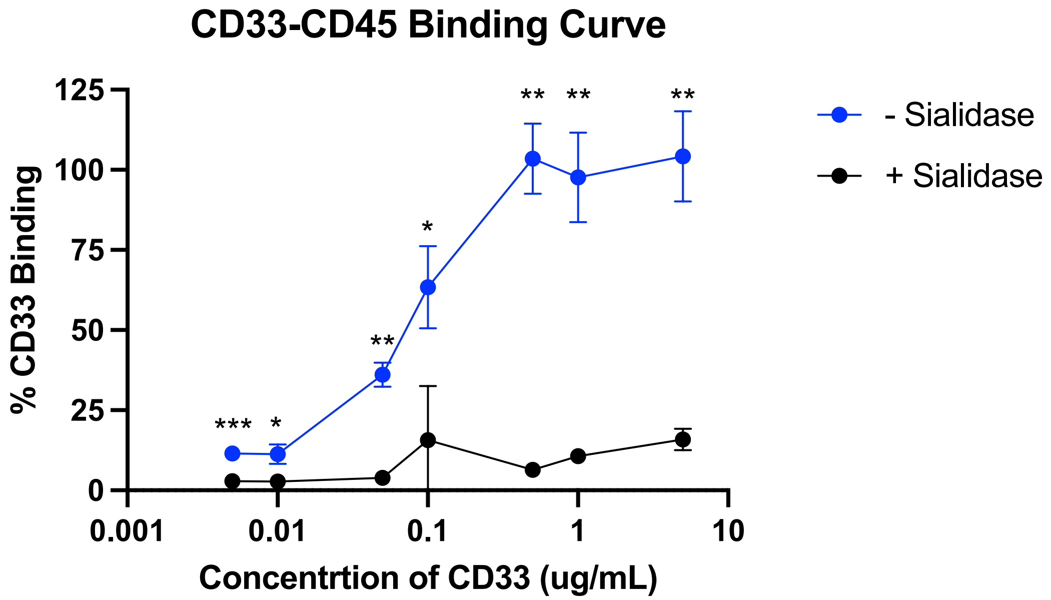


**Extended Data Figure 4. Binding of recombinant human CD33 to human CD45 is sialic acid dependent**. Treatment of recombinant human CD45 with 500mU/mL of neuraminidase (sialidase) inhibits binding to human CD33. Binding values were normalized by averaging the maximal absorbance values, which are represented as 100% binding. 0% binding was determined as the averaged absorbance values in the absence of recombinant human CD45. Each dot represents the normalized average of 4 technical replicates. Two-way ANOVA with Sidak's multiple comparisons test, *p<0.05, **p<0.01, ***p<0.01.

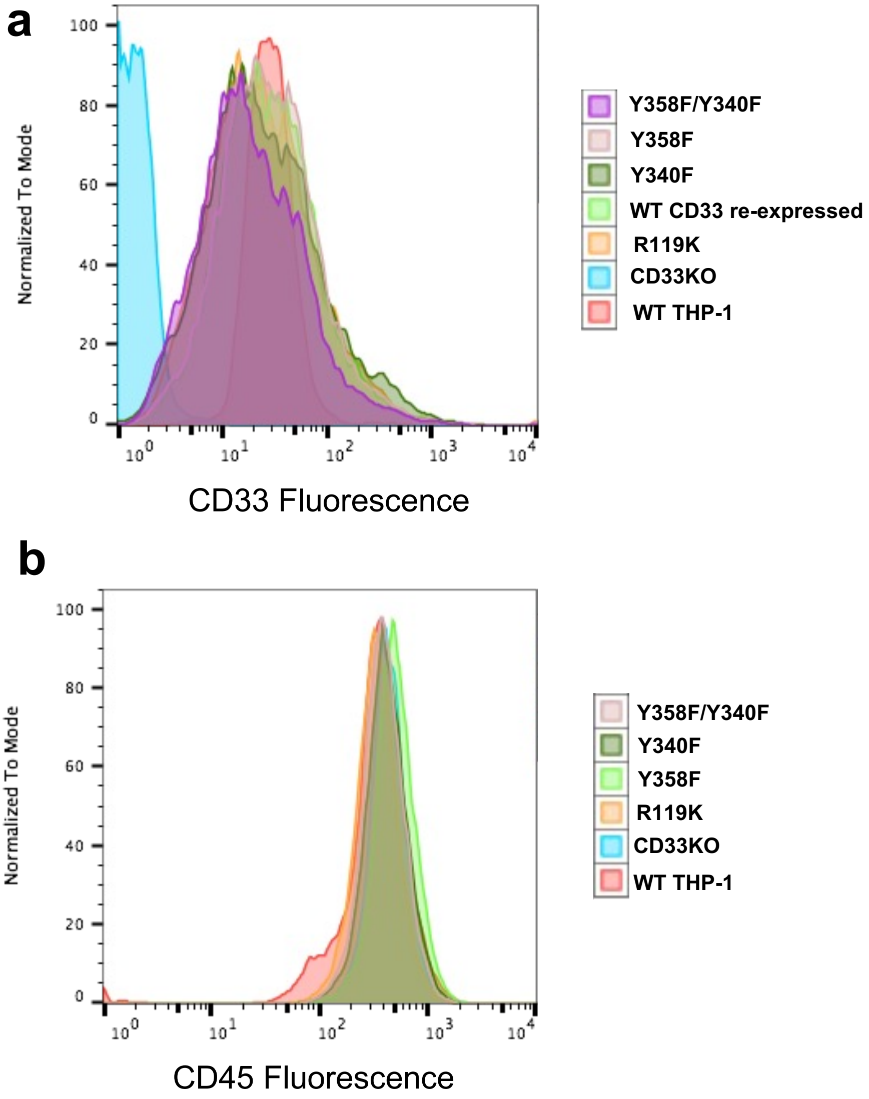


**Extended Data Figure 5. CD33 and CD45 protein expression in CD33 mutant THP-1 monocytes.** THP-1 monocyte lines were created expressing mutated CD33 at key functional sites within the protein. **a.** CD33 and **b.** CD45 expression in each line was assessed with flow cytometry.

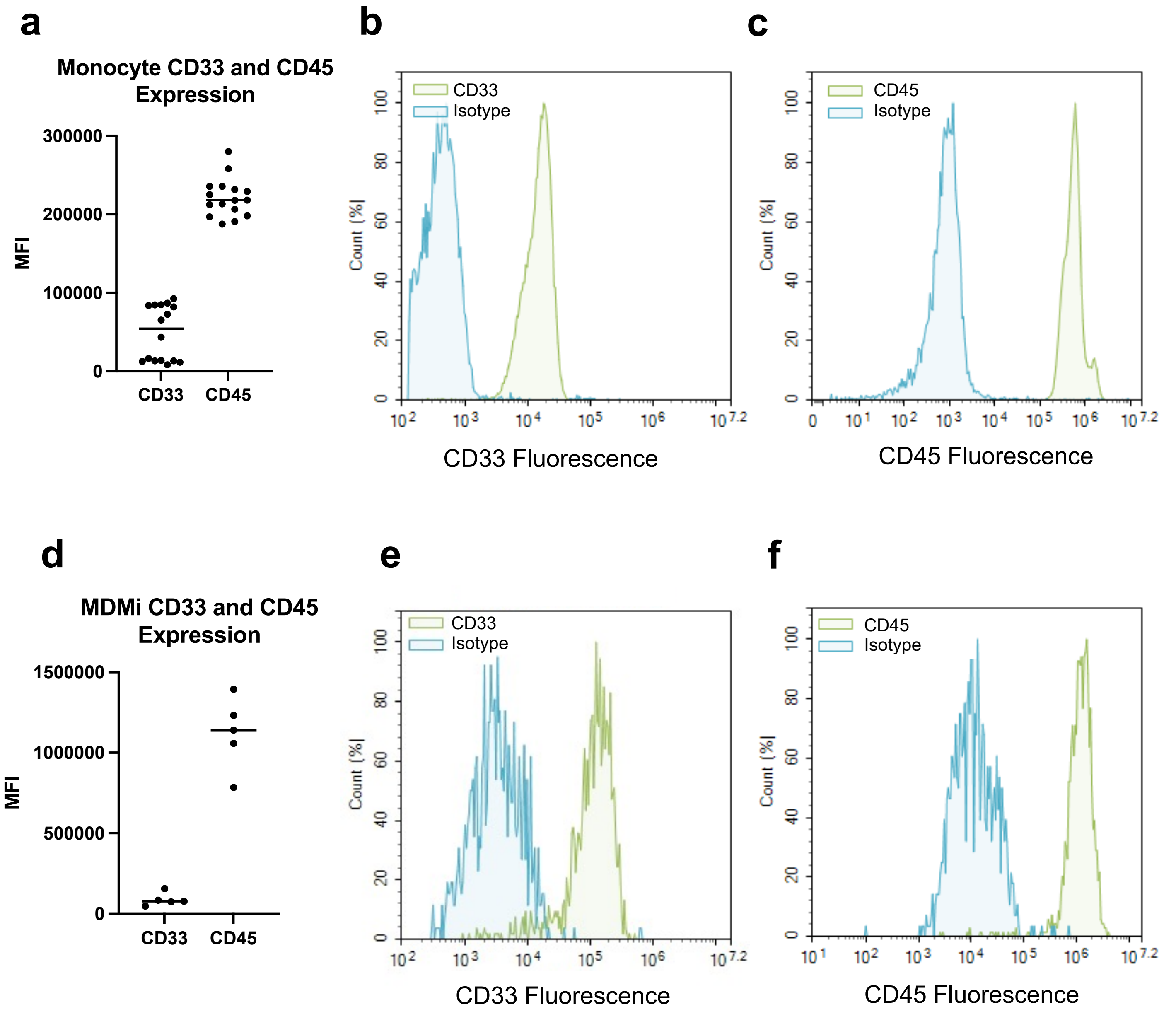


**Extended Data Figure 6. CD33 and CD45 protein expression in primary monocytes and monocyte-derived microglia (MDMi). a.** median fluorescence intensity of CD33 and CD45 expression in primary human monocytes. **b.** Representative flow cytometry histogram of CD33 staining versus isotype control in primary human monocytes. **c.** Representative flow cytometry histogram of CD45 staining versus isotype control in primary human monocytes. **d.** median fluorescence intensity of CD33 and CD45 expression in primary human MDMi. **e.** Representative flow cytometry histogram of CD33 staining versus isotype control in primary human MDMi. **f.** Representative flow cytometry histogram of CD45 staining versus isotype control in primary human MDMi.

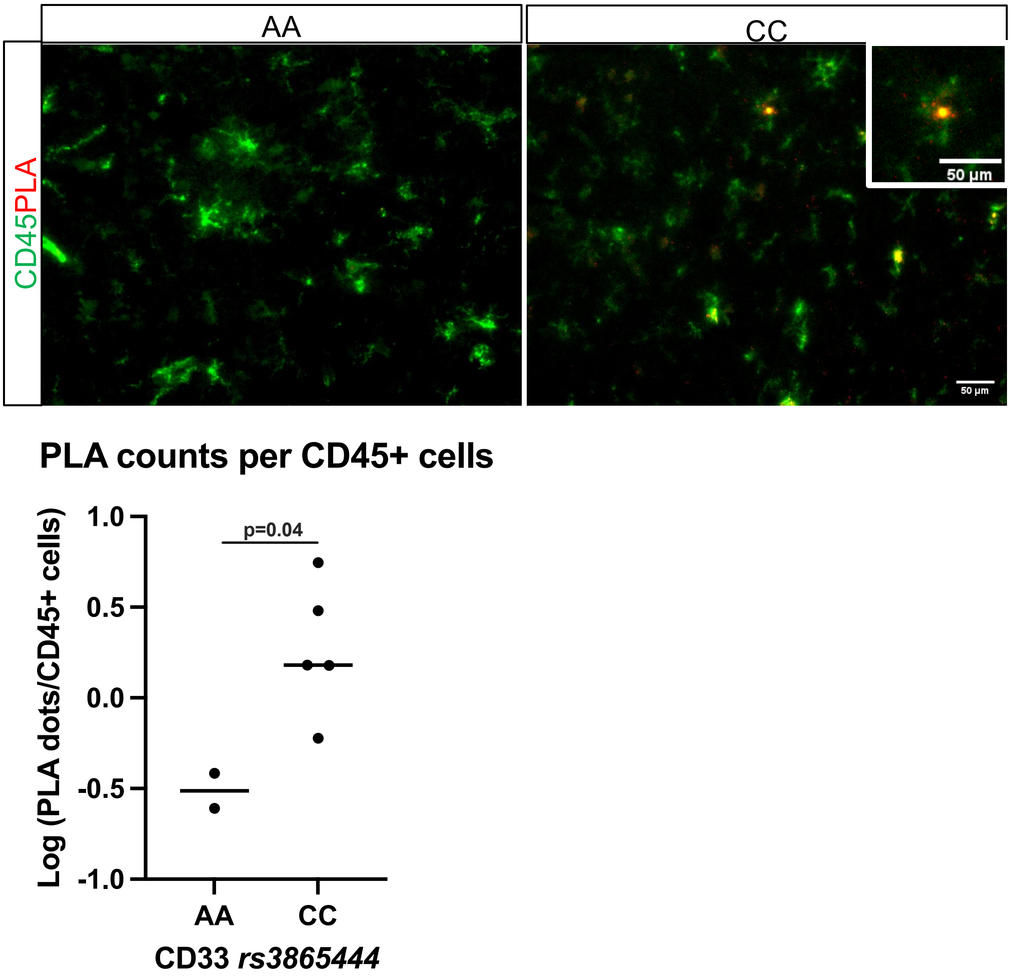


**Extended Data Figure 7: CD45 and CD33 Interact *in situ,* detected by proximity ligation assay**. Individuals with the CD33 homozygous protective genotype (N=2) have less CD33:CD45 interaction than individuals homozygous for the risk allele (N=5). Unpaired t-test of log transformation. Demographic and neuropathologic information for each individual is included in Supplementary Table 3.

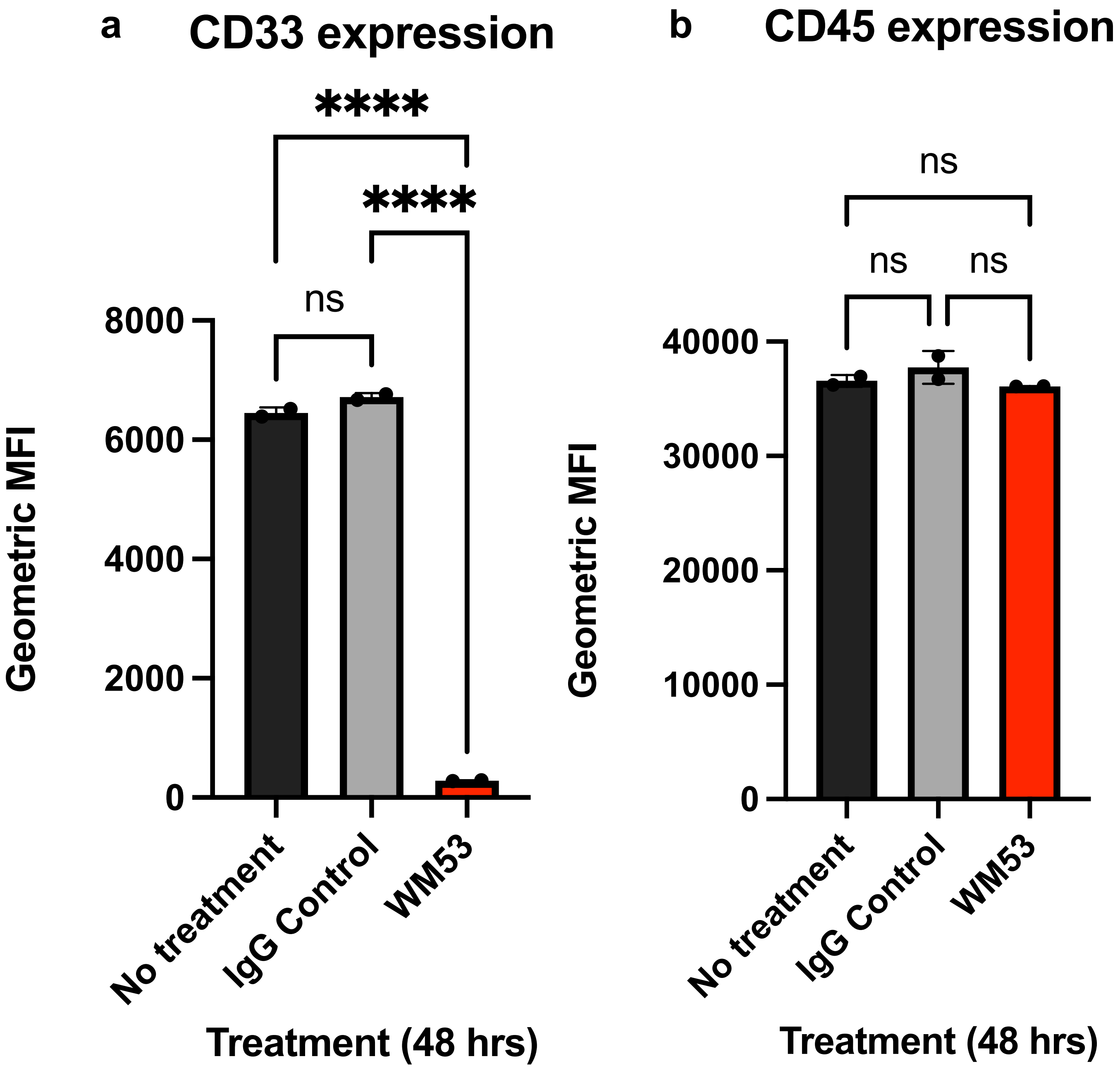


**Extended Data Figure 8. Treatment with the anti-CD33 antibody WM53 profoundly decreases surface CD33 expression without impacting CD45 expression.** THP-1 monocytes were treated with 10ug/mL of WM53 anti-CD33 antibody or IgG control at 37$^{o}$C for 48 hours. **a.**CD33 and **b.** CD45 surface protein expression were assessed with flow cytometry. Geometric mean florescence intensity (MF) is plotted.

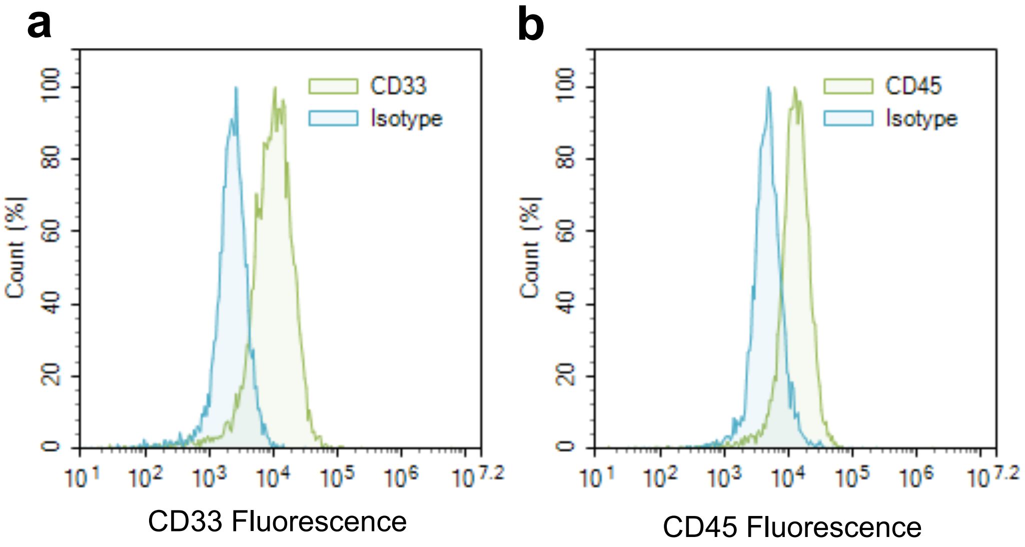


**Extended Data Figure 9. CD33 and CD45 protein expression in the HMC3 human microglial cell line.** **a.** Representative flow cytometry histogram of CD33 staining versus isotype control in HMC3 cells. **b.** Representative flow cytometry histogram of CD45 staining versus isotype control in HMC3s.

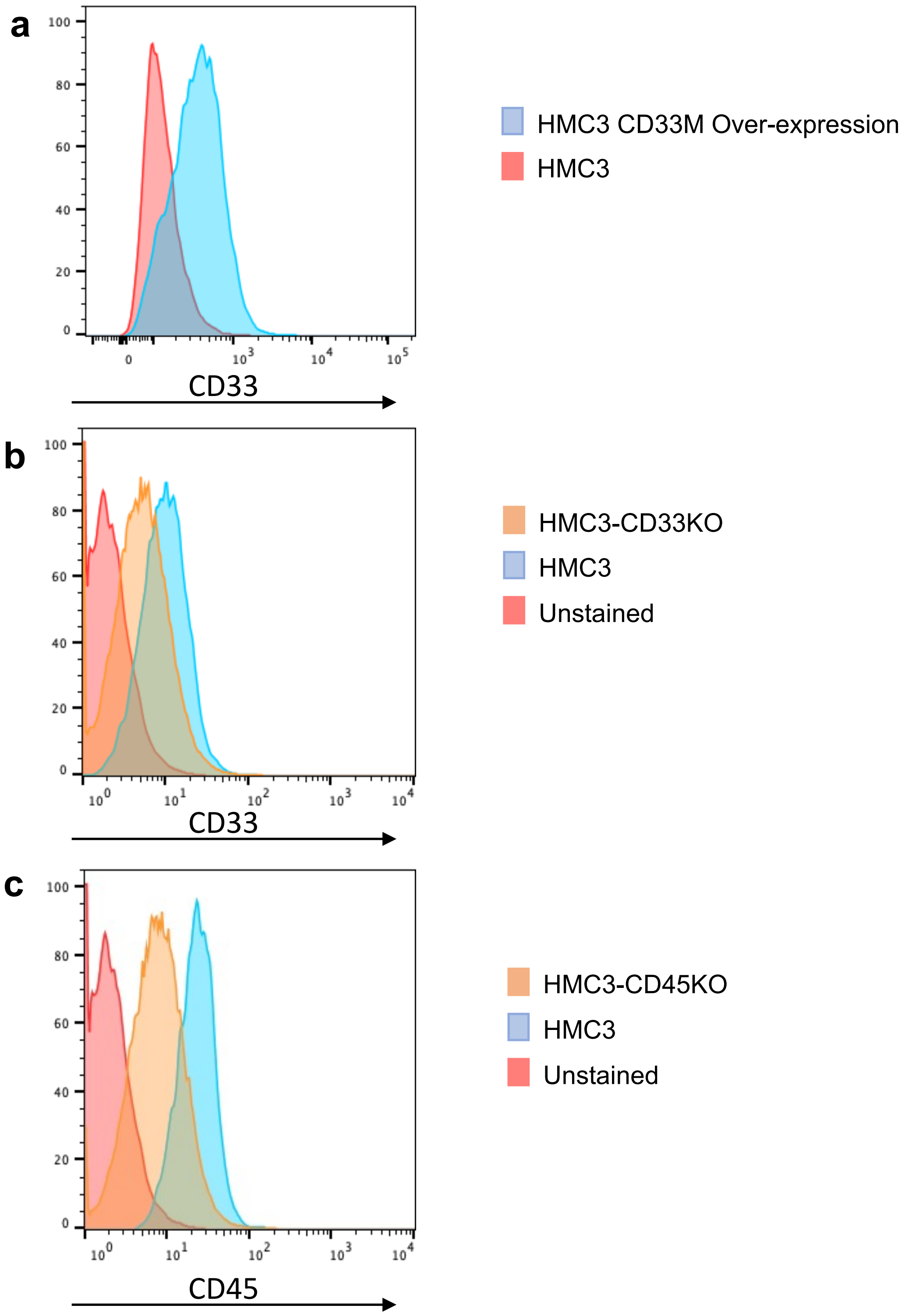


**Extended Data Figure 10: CD33 and CD45 expression in Knock-out and Overexpression HMC3 human microglial cell lines. a.** CD33 expression was assessed in WT HMC3 cells and HMC3s with over-expressed CD33M with flow cytometry. **b.** CD33 was knocked-out (KO) from the HMC3 cell line using Crispr/Cas9, and CD33 protein expression was assessed. **c.** CD45 was knocked-out from the HMC3 cell line using Crispr/Cas9, and CD45 protein expression was assessed with flow cytometry.

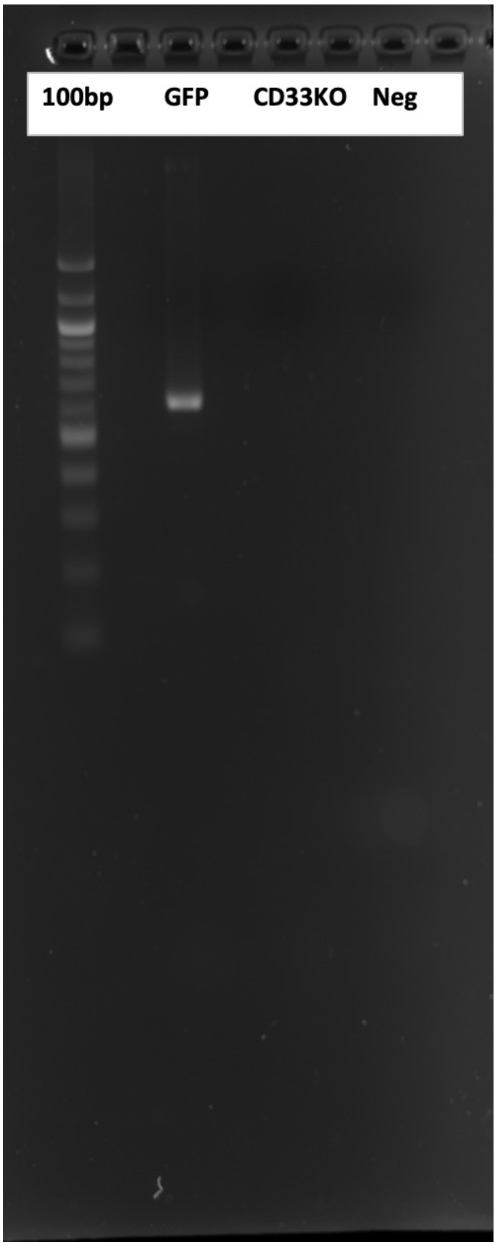


**Extended Data Figure 11: Confirmation of CD33 HMC3 Knockout (KO) lines.** CD33 was knocked out from the HMC3 cell line using Crispr/Cas9. Knockout was confirmed with agarose gel electrophoresis of PCR products (amplicons) amplified from genomic DNA at *CD33* exon 1 locus. DNA samples were run on a 1.5% agarose gel stained with ethidium bromide and visualized under UV light. Lane M: 100 bp DNA ladder (Thermo Fisher, SM0243), Lane 1: Amplicons generated from genomic HMC3 expressing GFP cells (~650 bp), Lane 2: Amplicons generated from genomic HMC3 CD33 knockout cells, Lane 3: negative control (no template).

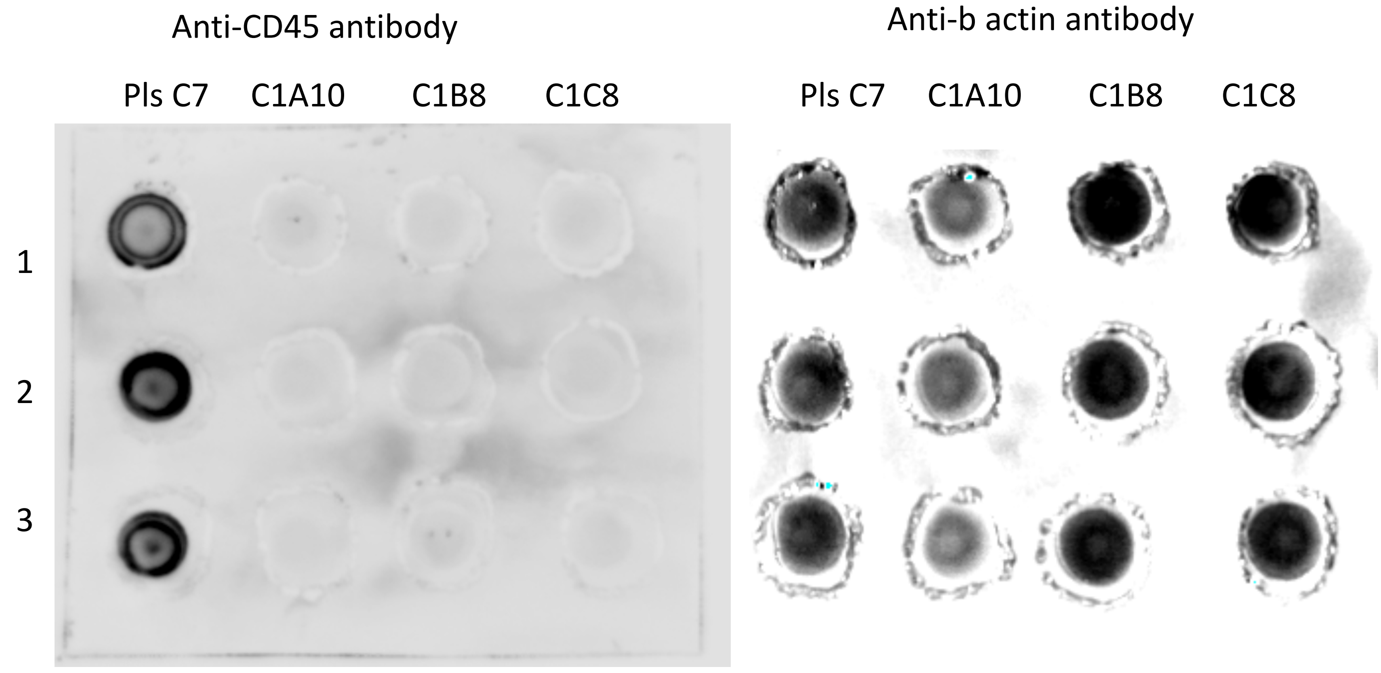


**Extended Data Figure 12: Confirmation of CD45 HMC3 Knockout (KO) line.** CD45 was knocked out from the HMC3 cell line using Crispr/Cas9. Knockout was confirmed by dot blot, performed in triplicate. PlsC7 served as a positive plasmid control. C1C8 represents the CD45KO clone used in subsequent experiments.
